# Targeting Non-Catalytic Sites of SRC Sensitizes the Efficacy of SRC Kinase Inhibitors in Solid Tumors

**DOI:** 10.1101/2025.02.15.638389

**Authors:** Jinmei Yu, Xusheng Wang, Pengyue Liu, Zhenhe Wang, Jianli Liu, Qianwen Zhang, Zewen Zhang, Tian Gan, Ning Xu, Jichao Zhou, Jiaojiao Yu, Hongyu Yuan, Xinchao Ban, Yuanzhen Liu, Xiaowei Zhang, Pingping Li, Bing Cui

## Abstract

The non-receptor tyrosine kinase SRC promotes the progression of hematologic malignancies and numerous solid tumors. Several SRC kinase inhibitors have been approved for the treatment of hematologic malignancies. However, drug resistance to sole ATP pocket-targeted kinase inhibition, coupled with low potency for solid tumors and insufficient selectivity, has limited the clinical application of SRC kinase inhibitors. The non-catalytic functions of kinases have provided crucial mechanisms underlying kinase inhibitor resistance, thereby presenting new opportunities for discovery SRC-targeted strategies. In this study, we discovered that upon abrogation of phosphorylation by SRC kinase inhibitors, non-catalytic SRC promotes the transcription of the oncogenes *TRIB3* and *SPC24* transcription by binding to their promoter sequence. Simultaneously, it interacts with TRIB3 and prevents their proteasome-mediated degradation by the E3 ligase CHIP, ultimately inducing resistance to SRC kinase inhibitors. The peptide TS1-2, blocking SRC/TRIB3 interaction in non-catalytic regions, inhibits the tumor progression of pancreatic ductal adenocarcinoma (PDAC), kidney renal clear cell carcinoma (KIRC), liver hepatocellular carcinoma (LIHC), and breast invasive carcinoma (BRCA), and enhances the efficacy of SRC kinase inhibitors in solid tumors. This study provides the insights of non-catalytic functions of SRC, validates the tumor-promoting function and mechanism of nuclear accumulation of SRC/TRIB3 following SRC kinase inhibitor treatment, identifies SRC/TRIB3 interaction as a potential non-catalytic target, and offers the candidate peptide TS1-2 as a potential combination strategy for overcoming SRC kinase inhibitor resistance.

**Significance:** The scaffolding functions of SRC reduced the efficacy of SRC kinase inhibitors, and targeting SRC/TRIB3 non-catalytic functions synergizes the therapeutic effects of SRC kinase inhibitors in SRC-dependent solid tumors.

**Graphical Abstract:** 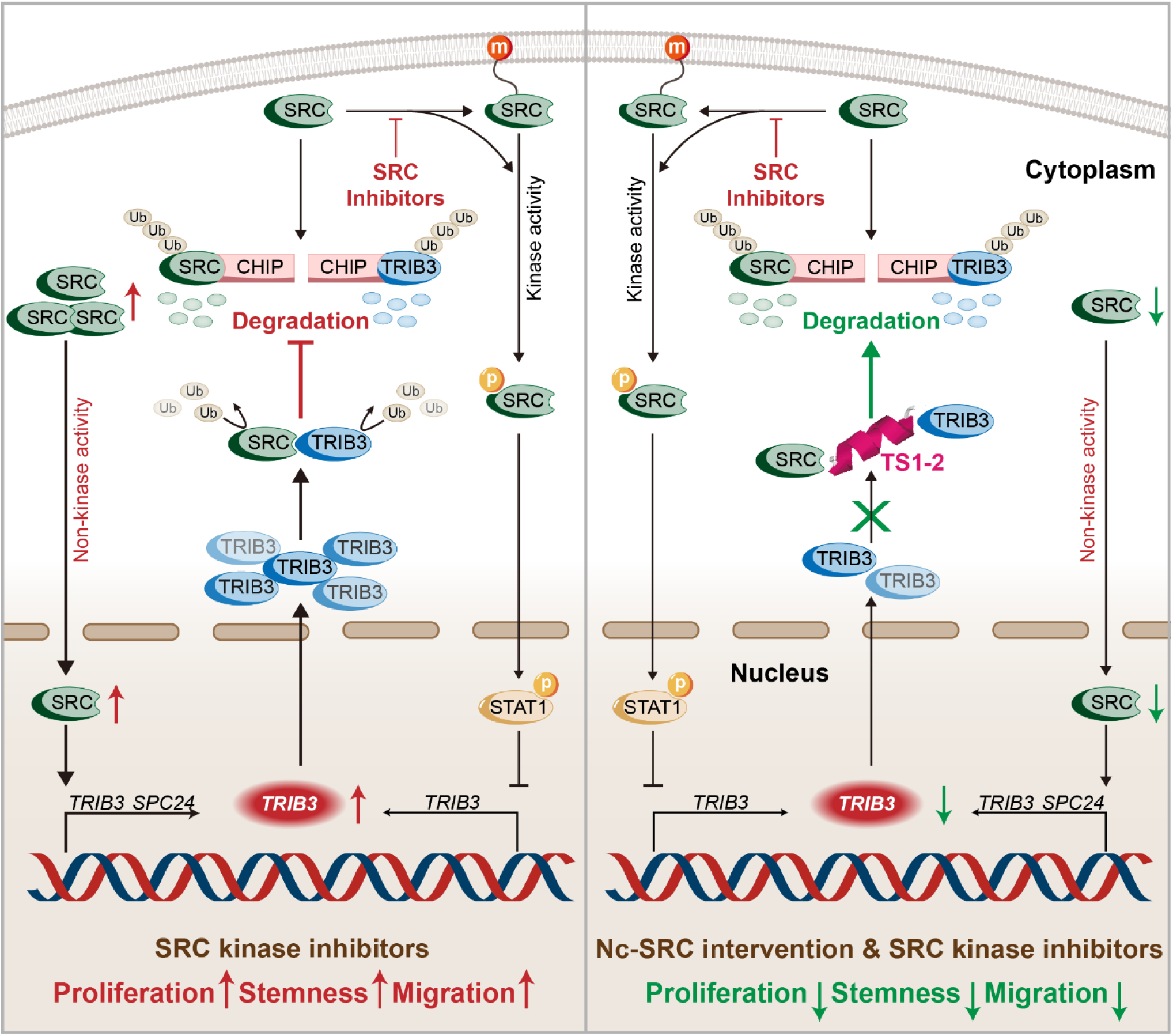

SRC kinase inhibitors block myristoylation-dependent membrane location and catalytic phosphorylation of SRC. Phosphorylated SRC activates STAT1 and thereby represses TRIB3 transcription. Accumulated non-catalytic SRC promotes *TRIB3*, *SPC24* transcription. Upregulated TIRB3 interacts with SRC to reduce CHIP-mediated ubiquitin–proteasomal degradation. The peptide TS1-2 disrupts TRIB3/SRC complex by targeting Non-kinase regions of SRC, consequently enhancing the therapeutic efficacy of SRC kinase inhibitors.

## INTRODUCTION

Protein kinases play pivotal roles in the post-translational modification of substrate proteins (1,2). They trigger conformational changes and subsequent biological activation of substrate proteins by transferring the terminal phosphate group of adenosine triphosphate (ATP) to specific tyrosine, threonine or serine residues (3). This phosphorylation process is facilitated by a highly conserved catalytic domain containing a canonical ATP-binding pocket. The human genome encodes approximately 518 protein kinases, which accounts approximately 2% of the protein-coding genome. Dysregulation of these kinases has been well-documented to be tightly associated with diverse human diseases, including malignancies, metabolic disorders, cardiovascular diseases, degenerative conditions, as well as immunological and infectious diseases (4). Globally, approximately 30% of pharmaceutical drug discovery efforts target the kinase superfamily (5). As of the latest updates the U.S. Food and Drug Administration (FDA) has approved 94 small-molecule kinase inhibitors for clinical applications (6,7). The majority of these approved kinase inhibitors exert their therapeutic effects via competitive binding to the ATP pockets of target kinases (8) or through a combination of allosteric and orthostatic modulation (9). However, the high structural similarity of ATP-binding pockets across different kinase families gives rise to off-target toxicity, a major factor contributing to the failure of kinase inhibitor-based clinical trial (10,11). Furthermore, acquired drug resistance also stands as another leading cause of poor clinical efficacy and even clinical trial failures, with point mutations within the ATP-binding pocket representing a major underlying mechanisms (12,13).

Recently, accumulating evidence demonstrates that the non-catalytic functions of kinases play crucial roles in diverse cellular processes through scaffolding functions. These include allosterically regulating other kinases via protein−protein interactions (PPIs), acting as scaffolds to assemble new signaling complexes, binding transcriptional factors(TFs) or directly interacting with DNA to modulate target genes expression (1,14–16). For example, kinase-inactive EGFR binds the mitochondrial pro-apoptotic protein PUMA in the cytoplasm, thereby inhibiting cancer cell apoptosis in glioblastoma (17). In breast cancer cells, kinase-inactive EGFR promotes starvation-induced tumor cell survival by facilitating autophagy initiation-via formation of a complex with LAPTM4B and Sec5. It also prevents autophagy-related cell death by stabilizing SGLT1 to maintain intracellular glucose levels (18,19). Structure and mutational analyses have further revealed that ERK2 binds to a GATE element (5’-CCCGGAGAGAATTGAAACTTAGGG-3’) in the target promoters via a surface pocket distinct from its ATP-binding pocket, thereby repressing IFNγ-induced genes expression (1,20). Notably, small-molecule modulators can either promote or disrupt kinase scaffolding functions through mechanisms such as orthostatic and allosteric conformation stabilization, targeted protein degradation, or direct protein-protein interaction (PPI) interception. Dr. Ding group discovered that degraders targeting kinases including protein kinase B (AKT3) (21), Anexelekto (AXL) (22), and tyrosine threonine kinase (TTK) (23), exhibit greater efficacy in ablating non-catalytic functions than in inhibiting catalytic activities. Despite the fact that non-catalytic functions of kinases have uncovered critical mechanisms underlying resistance to kinase inhibitors and presented new opportunities for kinase drug discovery, only a very limited number of modulators targeting these non-catalytic functions have been reported to date. Thus, exploring the molecular mechanism governing kinase the non-catalytic function represents a key strategy to overcome kinase inhibitor resistance and enhance the sensibility of cancer cells to kinase-targeted therapies.

SRC, a non-receptor tyrosine kinase and the first proto-oncogene identified by Bishop and Varmus in 1989, has been reported to interact with various receptors. These include receptor tyrosine kinases (EGFR, ErbB2, FGFR, c-Met, PDGFR), steroid hormone receptors (ER, PR) (24), integrins (αvβ3, β3), erythropoietin receptor (EPO) and G-protein-coupled receptors (AR, P2Y2) (25–27). These interactions synergistically activate multiple downstream signaling pathways, including the PI3K/AKT, STAT3, RAS, and FAK cascades (25,28), resulting in cell proliferation, survival, differentiation, adhesion, invasion, and angiogenesis (29,30). Consequently, SRC plays a critical role in the development and progression of human hematologic malignancies (31,32) as well as various solid tumors. Hematological malignancies driven by aberrant SRC activity include chronic myelogenous leukemia (CML) (33), acute myeloid leukemia (AML) (34), acute lymphoblastic leukemia (ALL) (35), while SRC-associated solid tumors comprise malignancies of the lung, colon, breast, ovarian, endometrial, skin, and head and neck (36,37). Several small-molecule ATP-competitive kinase inhibitors targeting SRC catalytic activity-including Dasatinib, Bosutinib, Saracatinib-as well as the non-ATP-competitive inhibitor Ponatinib, have been approved for the clinical treatment of hematologic malignancies (38–41). However, these agents have shown limited clinical efficacy against solid tumors, primarily due to the development of the drug resistance.

Accumulating evidence indicates that primary resistance to SRC inhibitors is largely driven by the loss of inhibitory functions for downstream signaling cascades. For instance, in Dasatinib-resistant head and neck squamous cell carcinoma (HNSCC) cells, the existence of the Src/c-Met interaction leads Dasatinib incapable of suppressing c-Met activity. Likewise, in Dasatinib-resistant pancreatic cancer cells, the phosphorylation levels of STAT3 and MAPK remain unaffected following Dasatinib treatment (42–44). In addition, the acquired resistance to SRC kinase inhibitors is the major cause of clinical trial failures, occurring primarily through the compensatory reactivation of downstream signaling pathways. For example, the aberrant activation of c-Met confers resistance to Dasatinib-induced apoptosis in gastric cancer (45). Sustained activation of the AKT/mTOR and MAPK pathways promotes Dasatinib resistance in thyroid cancer (46,47). In HER2-positive breast cancer cells, PAI-1 induces resistance to the Src inhibitor Saracatinib via the CCL5 axis (48). Prolonged Dasatinib treatment only causes transient inhibition of STAT3, followed by the restoration of STAT3 activity in a JAK-STAT3 binding-dependent manner in both HNSCCs (49) and non-small cell lung cancer (50). Notably, unlike Dasatinib which inhibits p-STAT3, silencing SRC paradoxically activates p-STAT3 in HNSCCs (49). Collectively, these findings strongly suggest that the non-catalytic function of SRC are critically involved in the development of resistance to SRC kinase inhibitors in solid tumors. However, the precisely regulatory mechanisms of SRC non-catalytic functions in this process remain largely unclear.

Given that esssential roles of non-catalytic functions of protein kinase, mainly by interacting with various partners, for inducing kinase inhibitor resistance, we postulated that pseudokinase TRIB3 interacted with SRC in contributes to the pathogenesis of several solid cancers via its non-catalytic functions. We studied the coordinative functions and mechanisms of the non-catalytic functions of SRC, stress sensor TRIB3, and E3 ubiquitin ligase CHIP in mediating SRC kinase inhibitor resistance, elucidated the implications of these findings, and provided potential therapeutic strategies using PPI blockers.

## Materials and Methods

### Cell culture

Human embryonic kidney cell line HEK293T, human liver hepatocellular carcinoma cell line HCCLM3, MHCC97H and HUH7, human kidney renal clear cell carcinoma cell line 769-P and 786-O, human breast cancer cell lines MDA-MB-231, MCF7 and murine breast cancer cell line 4T1 were obtained from National Infrastructure of Cell Line Resource, Peking Union Medical College (Beijing, China) and authenticated via short tandem repeats (STR) profiling. Human pancreatic ductal adenocarcinoma cell lines ASPC-1 and PANC-1 were acquired from American Type Culture Collection (ATCC, Manassas, VA, USA) and validated with the same STR for authentication.

### Plasmid construction

Various SRC truncation mutants were generated and constructed into the pCDH−Hyg-Myc expression vector via the BamHI and XbaI restriction sites. Additionally, the promoter fragments of *SPC24* and *BIRC5* were amplified and constructed into the pGL3-basic vector via KpnI and XhoI restriction enzyme digestion and ligation.

### Proximity ligation assay (PLA)

PLA was performed in accordance with the manufacturer’s protocol (Sigma, DUO96020). Anti-SRC (mouse; Abcam, ab231081) and anti-TRIB3 (rabbit; Abcam, ab75846) antibodies were used as primary antibodies. Fluorescence images were acquired with a laser scanning confocal imaging system (Olympus Microsystems).

### Animal Studies

BALB/c-nude mice were purchased from GemPharmatech Co., Ltd (Jiangsu, China) and housed in maximum barrier facilities with individually ventilated cages, sterilized food, and water. BALB/c mice were obtained from Beijing HFK Bioscience Co., Ltd. (Beijing, China) and maintained in the animal facility of the Institute of Materia Medica under specific-pathogen free (SPF) conditions.

See Supplementary Methods for detailed methods and references.

## RESULTS

### Non-catalytic SRC (Nc-SRC) mediates nuclear enrichment following SRC kinase inhibition

We queried and ranked the SRC protein and mRNA expression in 21 different solid tumors by utilizing Human Protein Atlas database (Supplementary Figure 1A and Table 1). We then queried The Cancer Genome Atlas (TCGA) database of different solid tumors with high-level SRC protein or mRNA expression by utilizing Kaplan-Meier plotter tools to interrogate if the impact of SRC expression was a significant predictor of improved or worsened overall survival (OS), recurrence-free survival (RFS) or disease-free survival (DFS) based on their relative SRC expression levels (Supplementary Table 1). Prostate, colon, stomach, urothelial, liver, pancreatic, renal, melanoma, breast cancer patients expressed relative high levels of *SRC* mRNA expression or SRC protein expression (Supplementary Table 1). In Pancreatic ductal adenocarcinoma (PDAC), Liver hepatocellular carcinoma (LIHC), Kidney renal papillary cell carcinoma (KIRP), Kidney renal clear cell carcinoma (KIRC), and Breast invasive carcinoma (BRCA) patients with tumors expressing *SRC* mRNA above the median level had a significantly shorter OS/DFS/RFS than patients with tumors expressing *SRC* below the median level (Supplementary Figure 1B). These SRC-dependent tumors were selected for further investigation, as these cancer types exhibit high expression of both SRC protein and/or mRNA, and high SRC mRNA expression is associated with poor survival prognosis (Supplementary Figure 1A-B and Supplementary Table 1).

Next, we focused on these SRC-dependent solid rumors to investigate the catalytic and non-catalytic role of SRC. First, the pancreatic cancer cell ASPC-1 sphere formation results indicated that silencing *SRC* synergized with 30nM SRC kinase inhibitor Dasatinib treatment (Supplementary Figure 1C). In order to understand the biological roles of Nc-SRC in SRC kinase inhibitor resistance, we conducted *in vitro* cell proliferation, cell migration and spheroid assays in SRC kinase inhibitor-resistant ASPC-1/BMS-R pancreatic cancer cells and HUH7/BMS-R liver cancer cells. It is interesting that silencing of SRC further inhibited cell proliferation, migration, and sphere formation in these SRC kinase inhibitor-resistant tumor cells (Supplementary Figure 1D-F). Moreover, the immunofluorescence results indicated that SRC was enriched in the nuclei of surviving subpopulations after 24-hour Dasatinib treatment in ASPC-1 cells (Supplementary Figure 1G). We constructed the phosphorylation-inactive mutant *SRC^Y419F^* mutants in HEK293 cells. With 24-hour Dasatinib treatment, SRC was mostly recruited to the cell membrane in the wild type SRC-expressing vector transfected cells, but enriched in the cell nucleus in the phosphorylation-inactive *SRC^Y419F^*-mutants transfected cells (Supplementary Figure 1H). Furthermore, western blotting results demonstrated that SRC was enriched in nuclear fractions of surviving HUH7 cell subpopulations following 24 h of Dasatinib treatment (Supplementary Figure 1I). Myristoylation has also been suggested to modulate the nuclear transport of SRC (53). Myristoylation assays revealed that both Dasatinib treatment and the *SRC^Y419F^*mutant suppressed the myristoylation of SRC (Supplementary Figure 2A), suggesting that Dasatinib promotes nuclear accumulation of SRC by inhibiting its myristoylation. These results indicate that Nc-SRC, especially its role in SRC nuclear accumulation, might play an important role in the survival of SRC kinase inhibitor-resistant cells. We hypothesized that Nc-SRC could directly regulate the transcription of certain genes to contribute to the survival of kinase inhibitor-resistant tumor cells.

### Nc-SRC promotes *SPC24* transcription via direct binding to the promoter sequence

To screen and identify the candidate Nc-SRC-regulated genes, we first analyzed the overlap of SRC target genes between the hTFtarget database and GSE224876 dataset. The latter contained differentially expressed genes (DEGs) identified from mammary epithelial cells isolated from PyMT transgenic mice, a well-established spontaneous breast cancer model, with mammary gland-specific knockout of SRC (SRC^−/−^). A total of 128 overlapping genes were identified. Notably, 13 upregulated oncogenes and 11 downregulated tumor suppressor genes (TSG) were screened out, based on their significant association with survival prognosis. This analysis utilized clinical data from TCGA database and the online Kaplan-Meier Plotter tool. Subsequently, mRNA analyses were performed following small interfering RNA (siRNA)-mediated SRC silencing and SRC inhibitor treatment. To further verify Nc-SRC target genes, additional analyses, including mRNA expression detection, promoter activity assays and Chromatin immunoprecipitation-PCR (ChIP-PCR) were conducted using the phosphorylation-inactive *SRC^Y419F^* mutants (Figure 1A).

**Figure 1.**
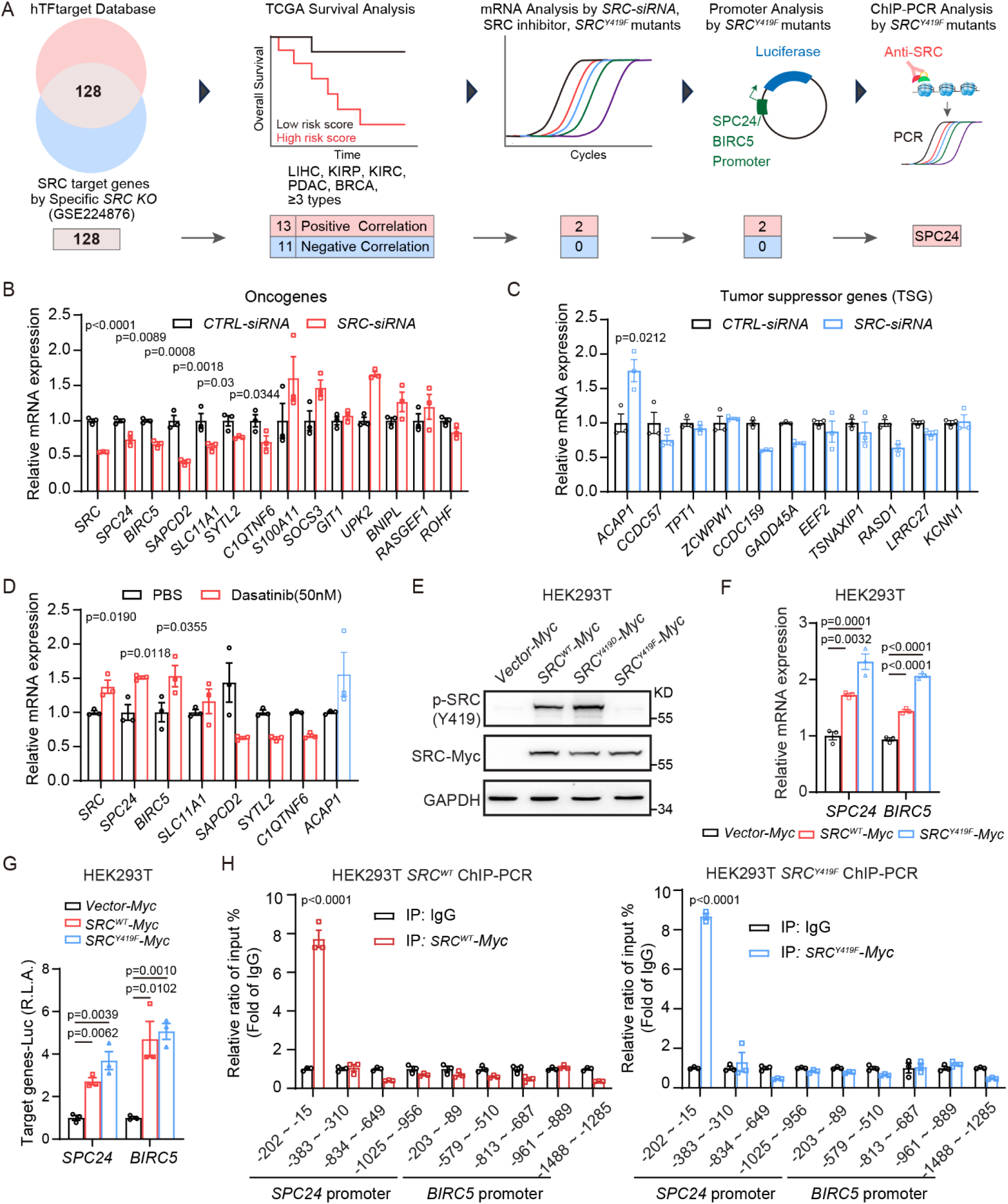
Non-catalytic SRC (Nc-SRC) promotes *SPC24* transcription via direct binding to the promoter sequence. **A,** Schematic diagrams of screening the target genes of Nc-SRC. First, SRC target genes were obtained from hTFtarget database, then 128 common SRC-targeted differentially expressed genes were selected based on GSE224876 dataset, with fold change ≥ 1.5 and *p* < 0.05. Second, overall survival analysis of LIHC, KIRP, KIRC, PDAC and BRCA patients from TCGA database were conducted with high- and low-risk groups by median cutoff of common SRC target gene expression. The selection criteria were set as at least 3 datasets with significant consistent trend. Third, 13 positive- and 11 negative-correlated SRC-targeted genes were selected to conduct following validation analysis including RT-qPCR assay by siRNA and SRC inhibitor, promoter assay and CHIP-PCR assay. **B-D,** Histograms indicating the relative mRNA amount of variety genes, as indicated at the bottom of each histogram, detected via RT-qPCR in triplicate samples of ASPC-1 cells transfected with either *CTRL-siRNA* or *SRC-siRNA* for 13 positive- (B) and 11 negative- (C) correlated SRC-target gene, or treated with either PBS or 50nM SRC inhibitor Dasatinib for 24h (D). **E,** Immunoblots of protein lysates of HEK293T transfected with a control vector, a *SRC^WT^*-expressing vector, a constitutively active mutants *SRC^Y419D^*-expressing vector or a kinase inactive mutants *SRC^Y419F^*-expressing vector as indicated at the top. **F-H,** The transcription of *SPC24* and *BIRC5* by Nc-SRC is validated in HEK293T cells transfected with different vectors by RT-qPCR (F) and the luciferase reporter assay (G). The binding regions is determined by the ChIP-PCR assay (H). n = 3 independent experiments in D-H, Data are shown as mean ± SEM; *P* > 0.05 is considered not significant (N.S.), \**P* < 0.05, \*\**P* < 0.01, \*\*\**P* < 0.001, compared with *CTRL-siRNA* or *CTRL-Vector* group.

Silencing SRC reduced the mRNA levels of six oncogenes and increased the mRNA level of one TSG in ASPC-1 cells, as verified by real-time PCR (Figure 1B-C). Furthermore, two of these oncogenes were enriched in surviving subpopulations of ASPC-1 cells following 24-h treatment with 50nM Dasatinib (Figure 1D). In HEK293T cells, overexpression of the kinase-inactive SRC mutant (*SRC^Y419F^*) (Figure 1E) enhanced the transcriptional activity of two oncogenes (*SPC24, BIRC5*). This was confirmed by real-time PCR (Figure 1F) and luciferase reporter assay (Figure 1G). Additionally, ChIP-PCR assays demonstrated that phosphorylation-inactive *SRC^Y419F^*directly bound to the promoter region of SPC24, but not that of BIRC5, in HEK293T cells (Figure 1H). Furthermore, silencing SRC decreased SPC24 expression in ASPC-1/BMS-R and HUH7/BMS-R cells (Supplementary Figure 2B). Notably, *SRC* silencing exerted a more potent inhibitory effect on sphere-forming capacity than *SPC24* silencing in ASPC-1/BMS-R and HUH7/BMS-R cells (Supplementary Figure 2C-D). Collectively, these results indicated that SRC promotes *SPC24* transcription via direct binding to its promoter region, in a catalytic activity-independent manner.

### The SRC/TRIB3 interaction is enriched in the nucleus after Dasatinib treatment

The Nuclear enrichment of SRC in cells expressing the phosphorylation-inactive *SRC^Y419F^*-mutants (Supplementary Figure 1H), together with the role of Nc-SRC in regulating gene transcription, prompted us to screen and identify key protein-protein interactions (PPIs) enriched in the nucleus of surviving cell subpopulations following SRC kinase inhibitor treatment. We then analyzed the overlapping upregulated genes across three SRC kinase inhibitor-related datasets: two datasets of renal carcinoma cells (GSE81235) and melanoma cells (GSE166617) treated with the SRC kinase inhibitor Dasatinib, and one dataset of hepatocellular carcinoma cells resistant to SRC kinase inhibitor Saracatinib (GSE129071). A total of 21 overlapping nuclear genes were obtained. Notably, 10 upregulated oncogenes were screened out, based on their significant association with overall survival prognosis in patients with LIHC, KIRP, KIRC, PDAC and BRCA from TCGA database (Figure 2A). Five of these 10 candidates interacted with SRC in surviving subpopulations following Dasatinib treatment, as identified via SRC CoIP-coupled mass spectrometry (Figure 2A-B and Supplementary Figure 3A). To further identify the key proteins that interacted with SRC and were upregulated in surviving subpopulations following Dasatinib treatment, we conducted mRNA and protein expression validation. Real-time PCR results showed that TRIB3 was the only gene whose mRNA expression was significantly upregulated in surviving subpopulations following Dasatinib treatment. (Figure 2C). A positive correlation between SRC and TRIB3 protein levels was observed in human PDAC, LIHC and KIRC cell lines (Figure 2D). Notably, patients with high TRIB3 expression exhibited poor survival in KIRC, LIHC, PDAC and BRCA cancers (Supplementary Figure 3B and Supplementary Table 2). Both the TRIB3 and SRC protein levels were markedly upregulated in surviving ASPC-1, 769-P, HUH7, and MCF7 cells following Dasatinib treatment (Figure 2E and Supplementary Figure 3C), as well as in Dasatinib-resistant ASPC-1/BMS-R and HUH7/BMS-R cells (Figure 2F). Additionally, Co-IP assays demonstrated that TRIB3 interacted with SRC in ASPC-1, 769-P, and HUH7 cells (Figure 2G). This physical interaction was further confirmed by Duolink in situ proximity ligation assay (PLA) in ASPC-1, 769-P, and HUH7 cells (Figure 2H) and immunofluorescence assay in human pancreatic cancer, liver cancer, kidney cancer and breast cancer (Supplementary Figure 3E). These results indicated that SRC/TRIB3 interaction might involve the functions of Nc-SRC in SRC kinase inhibitor resistant cells.

**Figure 2.**
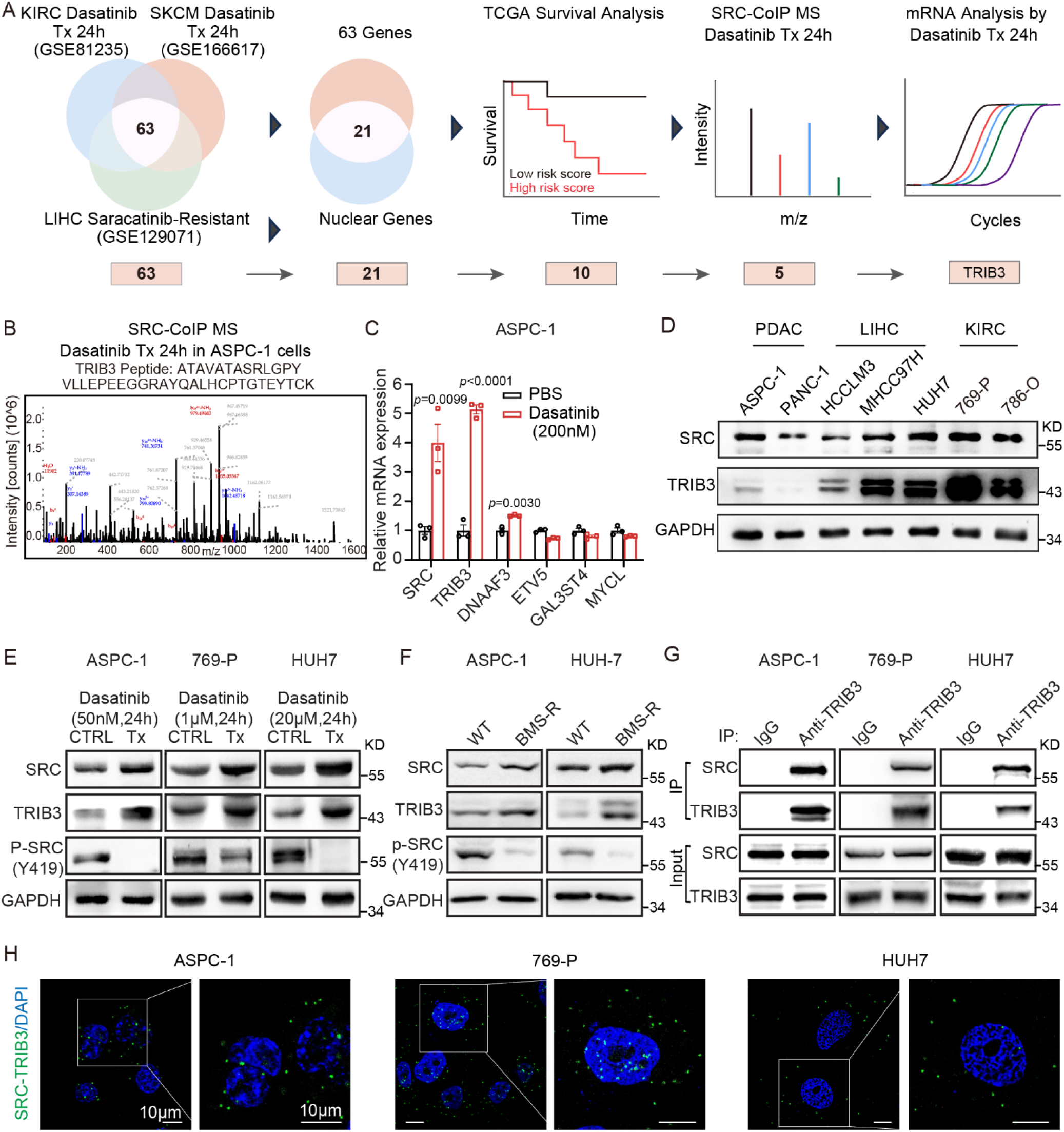
The TRIB3/SRC interaction is enriched in the nucleus after Dasatinib treatment. **A,** Schematic diagrams of screening SRC-related key protein interaction after SRC inhibitor Dasatinib treatment. First, 63 common SRC-up-regulated genes were obtained from tumor cells after SRC kinase inhibitor treatment (GSE81235, GSE166617) and SRC kinase inhibitor resistant cells (GSE129071), with old change ≥ 2 and p < 0.05. Then, 21 potential enriched nuclear genes were selected based on online Human Protein Atlas database. Second, overall survival analysis of LIHC, KIRP, KIRC, PDAC and BRCA patients from TCGA database were conducted with high- and low-risk groups by median cutoff of common SRC target gene expression. The selection criteria were set as at least 3 datasets with significant consistent trend. Third, 10 positive correlated genes were selected to conduct following validation analysis including Co-IP mass spectrometry and mRNA expression by Dasatinib treatment. **B,** SRC/TRIB3 interaction was identified by Co-IP mass spectrometry in ASPC-1 cells after 24 h Dasatinib treatment. **C,** Relative *SRC-*related nuclear genes mRNA expression in ASPC-1 cells after 24 h Dasatinib treatment. **D,** Representative Immunoblots probed for SRC, TRIB3 or GAPDH of lysates prepared from PDAC, LIHC, KIRC cells (as indicated on the top). **E-F,** Representative Immunoblots probed for TRIB3, SRC, p-SRC^Y419^ or GAPDH of lysates prepared from ASPC-1, 769-P and HUH7 cells after Dasatinib treatment for 24 h with different concentration (E, as indicated on the top), or from control and Dasatinib-resistant (BMS-R) ASPC-1/BMS-R and HUH7/BMS-Rcells (F, as indicated on the top). **G-H,** The TRIB3/SRC interaction in ASPC-1, 769-P and HUH7 cells validated by Co-IP (G) and Duolink PLA assay (H). n = 3 independent experiments in C-H, Data are shown as mean ± SEM; *P* > 0.05 is considered not significant (N.S.), \**P*<0.05, \*\**P*<0.01, \*\*\**P*<0.001, compared with *CTRL-siRNA*, PBS, *CTRL-Vector* or IgG group.

### SRC and TRIB3 synergistically promote the cell proliferation, migration and stemness in solid tumor cells

Next, we investigated the effects of TRIB3 and SRC on tumor progression. Combined siRNA-mediated silencing of TRIB3 and SRC exerted a more potent inhibitory effect on cell proliferation and migration compared with separate silencing of TRIB3 or SRC in ASPC-1, 769-P, HUH7 and MCF-7 cells (Figure 3A-3B and Supplementary Figure 4A). Silencing either TRIB3 or SRC reduced sphere formation ability. Notably, combined silencing of TRIB3 and SRC resulted in a more pronounced reduction compared with silencing either gene alone in the aforementioned four cell types (Figure 3C and Supplementary Figure 4B). These results indicate that SRC and TRIB3 synergistically promote the cell proliferation, migration and stemness of solid tumor cells. Subsequently, we investigated the cross-regulation between TRIB3 and SRC, as well as the underlying mechanism by which the SRC/TRIB3 interaction drives solid tumor progression.

**Figure 3.**
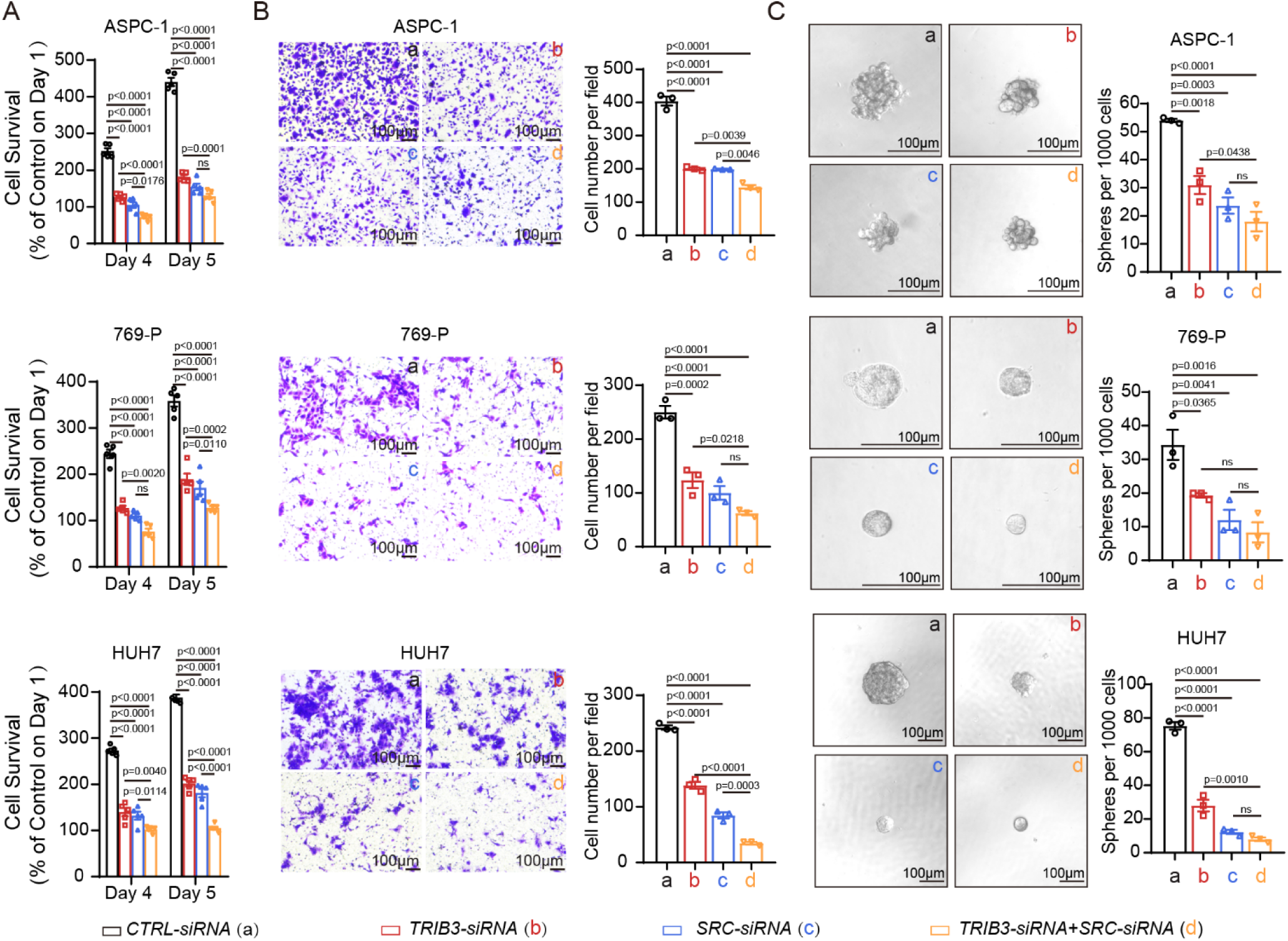
SRC and TRIB3 synergistically promote the cell proliferation, migration and stemness in solid tumor cells. Tumor proliferation (**A**), migration (**B**) and tumor sphere formation (**C**) analysis in ASPC-1, 769-P, and HUH7 cells transfected with *CTRL-siRNA* (a), *TIRB3-siRNA* (b), *SRC-siRNA* (c) or both *TIRB3-siRNA* and *SRC-siRNA* (d). The representative migration assay and tumor sphere formation photos are presented on the left of each panel (scale bar 100μM). n = 3∼5 independent experiments in A-C, Data are shown as mean ± SEM; *P*>0.05 is considered not significant (N.S.), \**P*<0.05, \*\**P*<0.01, \*\*\**P*<0.001, compared with *CTRL-siRNA*.

### SRC/TRIB3 interaction inhibits CHIP-mediated SRC ubiquitination and degradation

Firstly, silencing TRIB3 reduced SRC protein accumulation in ASPC-1, 769-P, HUH7 and MCF-7 cells (Figure 4A and Supplementary Figure 5A). We next investigated the mechanism by which TRIB3 upregulates SRC expression in solid tumor cells. Silencing TRIB3 did not significantly alter SRC mRNA levels in ASPC-1, 769-P, and HUH7 cells (Figure 4B). To evaluate whether TRIB3 regulates SRC protein stability, HEK293T cells were transfected with control (*CTRL)-HA* or *TRIB3-HA* vector for 24h, and followed by treatment with cycloheximide (CHX, 20 μM) for the indicated time. Notably, TRIB3 markedly prolonged the half-life of SRC by inhibiting its degradation (Figure 4C). Furthermore, TRIB3 silencing-induced reduction of SRC expression in ASPC-1 cells was abrogated by treatment with the proteasome inhibitor MG132 (10μM), but not by the autophagy inhibitor bafilomycin A1 (BAF, 200nM) (Figure 4D). Collectively, these data indicate that TRIB3 enhances SRC protein stability by interfering its degradation via the ubiquitin-proteasomal system (UPS).

**Figure 4.**
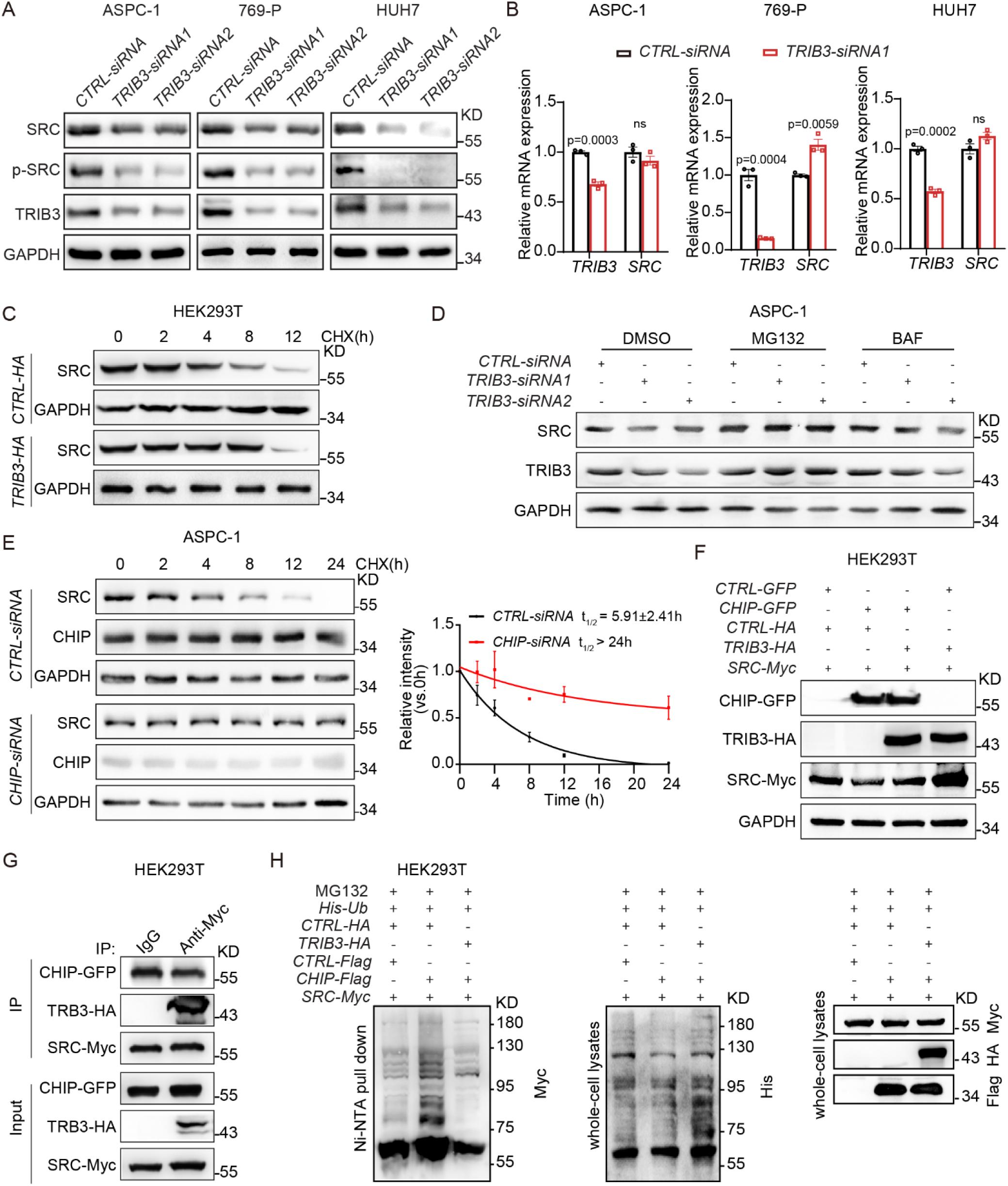
TRIB3/SRC interaction prevents the CHIP-mediated SRC ubiquitination and degradation. **A,** The representative immunoblots of protein lysates of SRC, p-SRC^Y419^ and TRIB3, as indicated on the left, in ASPC-1, 769-P and HUH7 cells transfected with *CTRL-siRNA*, *TRIB3-siRNA1* or *TIRB3-siRNA2*. **B,** Histogram showing the gene expression of *TRIB3* and *SRC* in ASPC-1, 769-P, and HUH7 cells transfected with *CTRL-siRNA*, *TRIB3-siRNA1*. **C,** The effect of TRIB3 overexpression on SRC degradation *in vitro*. HEK293T cells were transfected with a *TRIB3*-expressing vector (*TRIB3-HA*) or a control vector (*CTRL-HA*), and then incubated with 20 μM cycloheximide (CHX) for the indicated times. **D,** The effect of TRIB3 depletion on SRC degradation type *in vitro*. ASPC-1 cells were transfected with *CTRL-siRNA* or *TRIB3-siRNA* for 24 h, and then incubated with bafilomycin (200 nM) or MG132 (10 μM) for 8h. The indicated proteins were detected by immunoblotting. **E,** The effect of CHIP (gene name *STUB1*) depletion on SRC degradation *in vitro*. ASPC-1 cells were transfected with *CTRL-siRNA* or *CHIP-siRNA* for 24 h, then treated with CHX (20 μM) for the indicated time. The degradation curves and half-life are indicated on the right panel. **F,** TRIB3 reduced CHIP-mediated SRC degradation. HEK293T cells were transfected with the indicated plasmid for 24h, then cells extracts were blotted with an anti-GFP, anti-HA or anti-Myc Ab. **G,** TRIB3 inhibited the CHIP/SRC interaction. HEK293T cells were transfected with the indicated plasmid for 24 h. The cell extracts were immunoprecipitated with anti-SRC-Myc antibody and blotted with anti-CHIP-GFP or anti-TRIB3-HA antibody. **H,** The effect of TRIB3 on SRC ubiquitylation mediated by CHIP. HEK293T cells were transfected with the indicated plasmid. His-ubiquitin-conjugated proteins were pulled down with Ni-NTA-agarose from cell lysates. Total protein lysates and Ni-NTA-agarose eluates were immunoblotted and stained for FLAG, His, HA, and Myc. n = 3 independent experiments in A-G, Data are shown as mean ± SEM; *P*>0.05 is considered not significant (N.S.), \**P*<0.05, \*\**P*<0.01, \*\*\**P*<0.001, compared with *CTRL-siRNA*.

To identify the key E3 ubiquitin ligase mediating TRIB3-dependent inhibition of SRC degradation via the UPS, we first predicted SRC-interacting E3 ligases using the Unibrowser 2.0 database. Seven known SRC-targeting E3 ligases were selected for further investigation (Supplementary Figure 5B). To screen E3 ligases that interact with SRC, we analyzed the protein-protein interaction data from the SRC CoIP-coupled mass spectrometry assays. This analysis revealed that the interaction between SRC and E3 ligase FBXL7 or CHIP (Gene Name: *STUB1*) was abrogated following Dasatinib treatment (Supplementary Figure 5C-D). Previous studies have shown that FBXL7 mediates the ubiquitination and proteasome degradation of active SRC following its phosphorylation at Ser104, and that FBXL7 interacts with active SRC (i.e., p-SRC (Y419)) but not inactive SRC (i.e., p-SRC (Y530)). FBXL7 overexpression reduces endogenous p-SRC (Y419) levels, with a modest effect on total SRC but no impact on p-SRC (Y530) (54). These findings indicate that FBXL7 cannot mediate the degradation of inactive SRC. In contrast, Co-IP assays demonstrated that CHIP interacted with both active p-SRC (Y419) and inactive p-SRC (Y530) (Supplementary Figure 5E-F).

Given its ability to bind both SRC isoforms, CHIP was selected for subsequent studies. Silencing CHIP prolonged the half-life of SRC degradation from 5.9 h to >24 h (Figure 4E). Interestingly, silencing CHIP also extended the half-life of TRIB3 degradation from 1.84 h to >8 h (Supplementary Figure 5G). In HEK293T cells, CHIP overexpression decreased SRC protein levels, while concurrent overexpression of TRIB3 reversed this effect (Figure 4F). Furthermore, the TRIB3/SRC interaction blocked the CHIP/SRC interaction in HEK293T cells (Figure 4G). His-ubiquitin pull-down assay showed that ectopic CHIP expression enhanced SRC ubiquitination, whereas simultaneous overexpression of TRIB3 attenuated CHIP-mediated SRC ubiquitination in HEK293T cells (Figure 4H). Collectively, these data indicate that TRIB3 inhibits CHIP-mediated ubiquitin-proteasomal degradation of SRC.

### p- SRC potentiates p-STAT1-mediated repression of TRIB3 transcription, while Nc-SRC upregulates TRIB3 transcription

To further explore the role of SRC in regulating TRIB3 protein expression, we found that siRNA-mediated SRC silencing increased TRIB3 accumulation in ASPC-1, 769-P and HUH7 cells (Figure 5A). Notably, overexpression of the phosphorylation-inactive mutant *SRC^Y419F^* upregulated TRIB3 protein expression, while wild-type *SRC* reduced the TRIB3 protein levels in HEK293T cells (Figure 5B). Consistent with this, *SRC^Y419F^* overexpression enhanced *TRIB3* mRNA levels and promoter-driven luciferase reporter activity, whereas wild-type *SRC* overexpression exerted the opposite effects in HEK293T cells (Figure 5C-5D). Chromatin immunoprecipitation (ChIP) assays showed that both SRC^WT^ and SRC^Y419F^ proteins directly bound the *TRIB3* promoter region (Figure 5E). Collectively, these results indicate that p-SRC inhibits *TRIB3* transcription, while Nc-SRC promotes TRIB3 transcription via direct binding to the *TRIB3* promoter.

**Figure 5.**
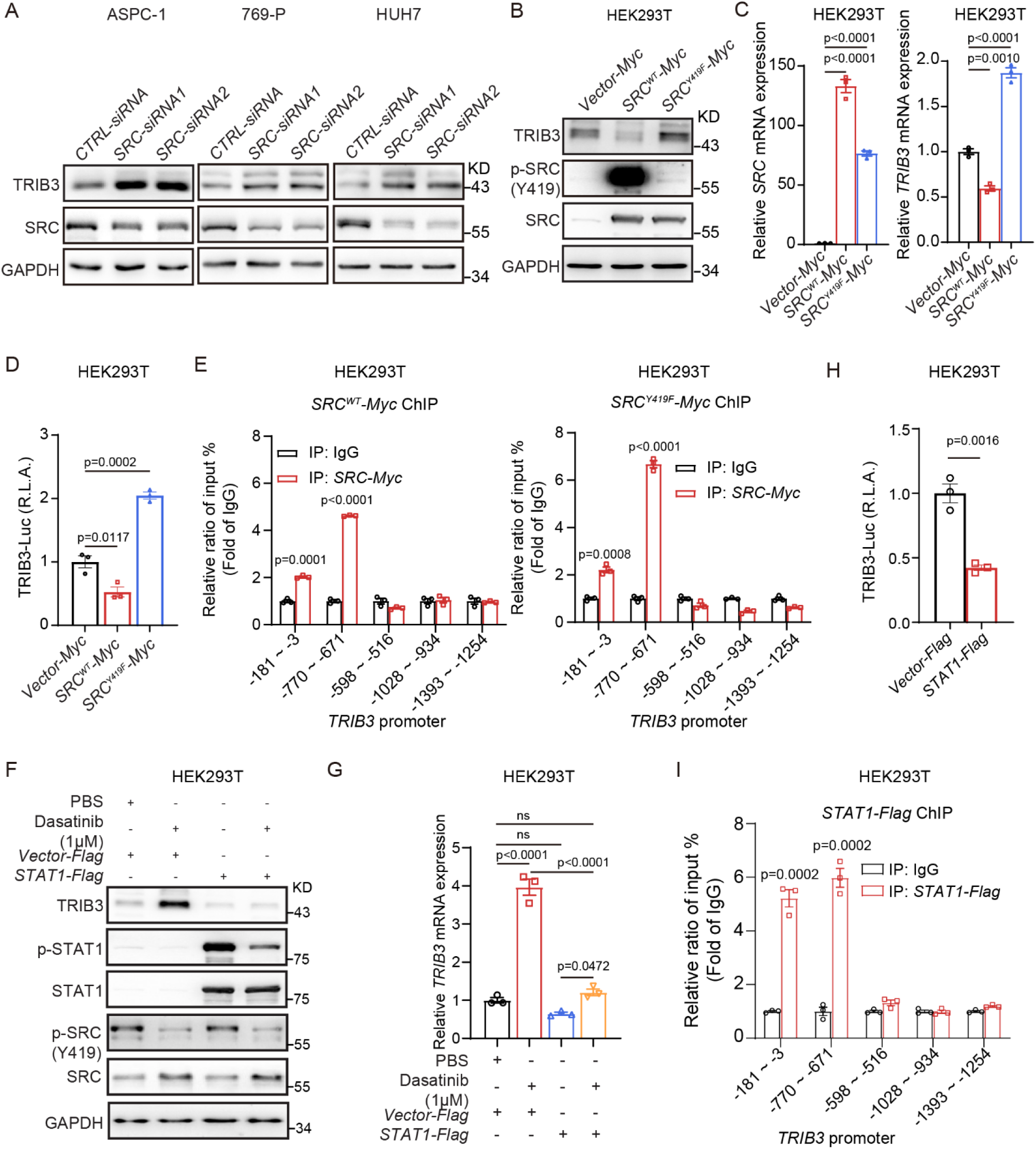
p-SRC^Y419^ inhibits but Nc-SRC promotes *TRIB3* transcription. **A,** Immunoblots of protein lysates, as indicated on the top, transfected with *CTRL-siRNA* or *SRC-siRNA* in ASPC-1, 769-P, and HUH7 cells. **B-D,** Immunoblots of protein lysates (B), RT-qPCR of cell lysates (C) and promoter-luciferase reporter assay of live cells (D), as indicated on the top or bottom, transfected with a control vector, a *SRCs^WT^*-expressing vector or a *SRC^Y419F^*-expressing vector in HEK293T cells. **E,** SRC is recruited to the *TRIB3* promoter regions. ChIP analysis was used to monitor the binding of SRC to the *TRIB3* promoter in HEK293T cells expressing a *SRC^WT^-Myc* vector (left) or *SRC^Y419F^-Myc* vector (right). **F-H,** Immunoblots of protein lysates (F), RT-qPCR of cell lysates (G) and promoter-luciferase reporter assay of live cells (H), as indicated on the top or bottom, transfected with a control vector, or a *STAT1*-expressing vector for 36 h in HEK293T cells with or without SRC inhibitor Dasatinib treatment for 24 h. **I,** STAT1 is recruited to the *TRIB3* promoter regions. ChIP analysis was used to monitor the binding of STAT1 to the *TRIB3* promoter in HEK293T cells transfected with *STAT1-Flag* vector. Data represent at least three separate experiments and are shown as the mean ± SEM., *P*>0.05 is considered not significant (N.S.), \**P*<0.05, \*\**P*<0.01, \*\*\**P*<0.001, compared with *CTRL-siRNA*, *CTRL-Vector* or IgG group.

To explore the mechanism by which p-SRC inhibits *TRIB3* transcription, we noted that nuclear SRC can enhance the tyrosine phosphorylation, DNA binding capacity, and transcriptional activity of transcription factors (e.g., STAT1, STAT3 and p300) via protein-protein interaction (55,56). We further predicted the TRIB3-targeting transcription factors using the online hTFtarget and JASPAR databases, and integrated these data with SRC-interacting proteins identified from the NCBI database and SRC CoIP-coupled mass spectrometry assays. This analysis yielded 13 candidate transcription factors (Supplementary Figure 6A-B), among which six (STAT1, STAT3, AR, STAT5B, ETS1, ETS2) are known substrates of phosphorylation-active SRC (Supplementary Figure 6C). Given our previous finding that TRIB3 inhibits *STAT1* transcription (57), we then focused on the cross-regulation of the p-SRC/p-STAT1/TRIB3 axis.

Dasatinib treatment markedly reduced p-SRC and p-STAT1 protein levels, while increasing TRIB3 expression in these cells (Figure 5F). Furthermore, STAT1 overexpression abrogated the Dasatinib-induced upregulation of TRIB3 protein (Figure 5F) and mRNA levels (Figure 5G), as well as the enhancement of TRIB3 promoter-driven luciferase activity (Figure 5H). ChIP-PCR assays confirmed that STAT1 directly bound to the TRIB3 promoter sequence in HEK293T cells (Figure 5I). These results indicated that p-SRC phosphorylates STAT1, and p-STAT1 inhibits *TRIB3* transcription through direct binding to its promoter; TRIB3, in turn, inhibits *STAT1* transcription, forming a negative feedback loop. Considering the fact that SRC promotes *SPC24* transcription in a catalytic activity-independent manner, we further demonstrated that combined silencing of *TRIB3* and *SPC24* produced a comparable inhibitory effect on tumor sphere formation relative to single knockdown of *SRC* in Dasatinib-resistant ASPC-1/BMS-R and HUH7/BMS-R cells (Supplementary Figure 6D). Collectively, these results suggest that SPC24 and TRIB3 synergistically mediate the non-catalytic function of SRC in Dasatinib-resistant cells.

### Targeting the SRC/TRIB3 interaction reduces solid tumor progression

We next mapped the key region of SRC mediating its interaction with TRIB3 via Co-IP assays using Myc-tagged *SRC* deletion mutants and HA-tagged *TRIB3*. TRIB3 specifically interacted with the 439-469 amino acid (aa) region within the kinase domain (Figure 6A-E). According to the UniProt database, this interaction region is distinct from the SRC ATP-binding site, which spans residues 276-284 aa, and includes residue 298 aa. To validate the biological function of the SRC/TRIB3 interaction in regulating of SRC expression and tumor progression, we sought to identify a short α-helical peptide derived from the 439-469 aa region of the SRC kinase domain that could inhibit the SRC/TRIB3 interaction. Peptide S1-2 (WSFGILLTELTTKG), a mimic of 448–462 aa α-helical region within the SRC kinase domain, exhibited specific binding affinity for TRIB3, as detected by ForteBio Bio-Layer Interferometry (BLI) (Figure 6F).

**Figure 6.**
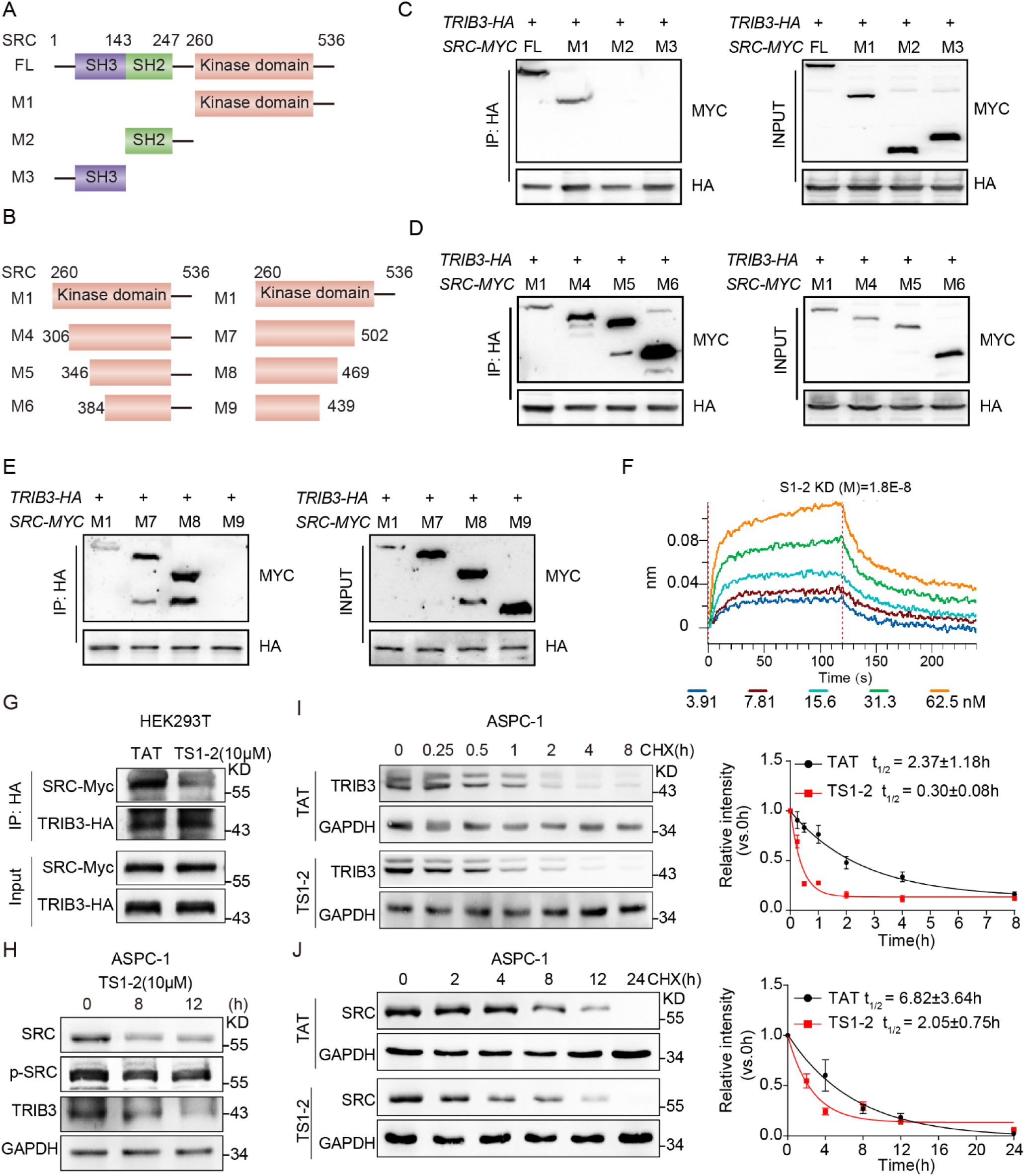
TS1-2 disturbs the SRC/TRIB3 interaction and downregulates the expression of SRC and TRIB3. **A-B,** Mapping SRC regions binding to TRIB3. Schematic diagram of full-length SRC and deletion mutants. **C-E**, HEK293T cells were co-transfected with the indicated *SRC-Myc* and *TRIB3-HA* constructs. Cell extracts were immunoprecipitated with anti-HA antibody. **F,** The kinetic interaction of α-helical peptide S1-2 and TRIB3 protein was determined by BLI analyses. **G,** TS1-2 interrupted the SRC/TRIB3 interaction. HEK293T cells were treated with 10 μM TAT or TS1-2 for 12 h. Cell extracts were immunoprecipitated with anti-HA antibody and blotted with an anti-TRIB3-HA or anti-SRC-Myc Ab. **H,** TS1-2 reduced both TRIB3 and SRC expression. ASPC-1 cells were treated with the TS1-2 peptide for different amounts of times. The indicated proteins were detected with immunoblotting. **I-J,** TS1-2 enhanced TRIB3 degradation (I) and SRC degradation (J). ASPC-1 cells were treated with 20 μM CHX and the indicated peptides for different amounts of times. The indicated proteins were detected with immunoblotting. Data represent at least three separate experiments and are shown as the mean ± SEM.

To determine whether S1-2 disrupts the SRC/TRIB3 interaction in solid tumor cells, we generated the cell-permeable fusion peptide TS1-2 by linking S1-2 to the TAT peptide (GRKKRRQRRR) (58) via a glycine–glycine (GG) linker. TS1-2 effectively abrogated the SRC/TRIB3 interaction in HEK293T cells (Figure 6G). Notably, TS1-2 simultaneously reduced the protein expression levels of both TRIB3 and SRC (Figure 6H). Furthermore, in ASPC-1 cells, TS1-2 significantly accelerated the degradation of TRIB3 and SRC, shortening their protein half-lives from 2.3 h to 0.3 h and from 6.8 h to 2 h, respectively (Figure 6I-J). These results indicate that the SRC/TRIB3 interaction not only stabilizes SRC proteins, but also blocks TRIB3 protein degradation, with the underlying mechanism warranting further investigation.

Since the SRC/TRIB3 interaction was found to be enriched in the nuclei of surviving cells following treatment with the SRC kinase inhibitor Dasatinib, we investigated the therapeutic effect of combining TS1-2 with Dasatinib. Results showed that compared with Dasatinib alone, TS1-2 monotherapy significantly suppressed cell proliferation (Figure 7A and Supplementary Figure 7A), migration (Figure 7B), and tumor spheres formation (Figure 7C and Supplementary Figure 7B). The combination of TS1-2 and Dasatinib exerted robust synergistic inhibitory effects in ASPC-1, 769-P and MCF-7 cells. In HUH7 cells, TS1-2 and Dasatinib monotherapies exhibited comparable inhibitory effects on cell proliferation, migration, and tumor spheres formation, whereas their combination still elicited marked synergistic effects (Figure 7A-C). Collectively, these results demonstrate that TS1-2 exerts a more potent inhibitory effect than Dasatinib by concurrently downregulating SRC and TRIB3 expression, and it exhibits robust synergistic antitumor effects with Dasatinib.

**Figure 7.**
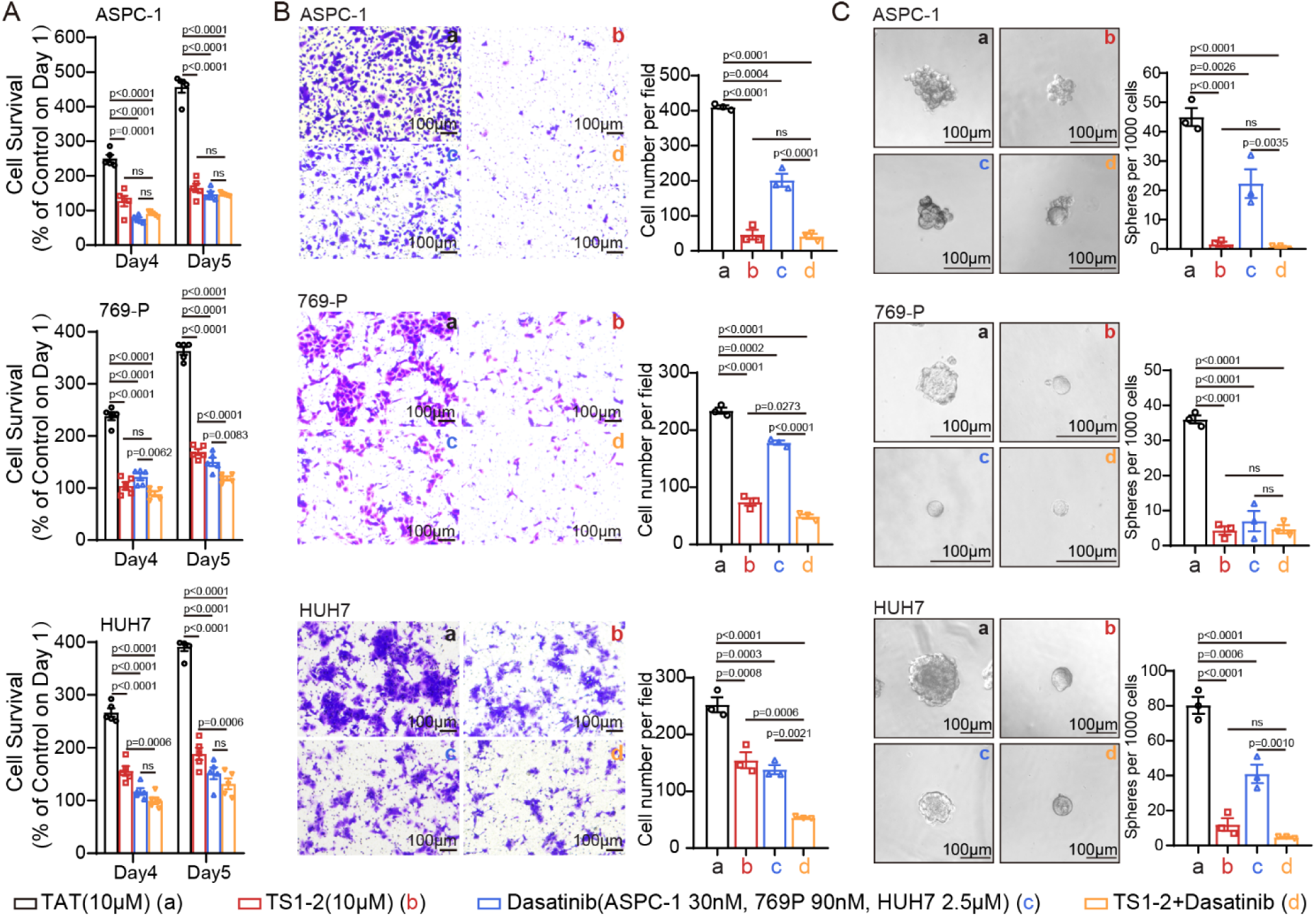
TS1-2 inhibits the tumor cell proliferation, migration, and stemness. Tumor proliferation **(A),** migration **(B)** and tumor sphere formation **(C)** analysis in ASPC-1, 769-P, and HUH7 cells treated with TAT (a), TS1-2 (b), Dasatinib (c) or both TS1-2 and Dasatinib (d). The representative migration assay and tumor sphere formation photos are presented on the left of each panel (scale bar 100μM). n = 3∼5 independent experiments in A-C, Data are shown as mean ± SEM; P>0.05 is considered not significant (N.S.), \**P*<0.05, \*\**P*<0.01, \*\*\**P*<0.001.

Combination treatment with TS1-2 and Dasatinib significantly reduced tumor growth and tumor weight, showing robust synergistic antitumor effects in ASPC-1 xenograft models in BALB/c nude mice (Figure 8A-D) and 4T1 xenograft models in BALB/c mice (Figure 8F-I). In tumor tissues derived from ASPC-1 and 4T1 xenografts, TS1-2 significantly downregulated the protein expression of SRC, TRIB3 and SPC24 (Figure 8E and 8J). Collectively, these data confirm that targeting the SRC /TRIB3 interaction impairs the cancer stemness in solid tumors and exerts potent antitumor efficacy by promoting the degradation of both SRC and TRIB3, which consequently suppresses SPC24 expression in pancreatic, kidney, liver, and breast cancers.

**Figure 8.**
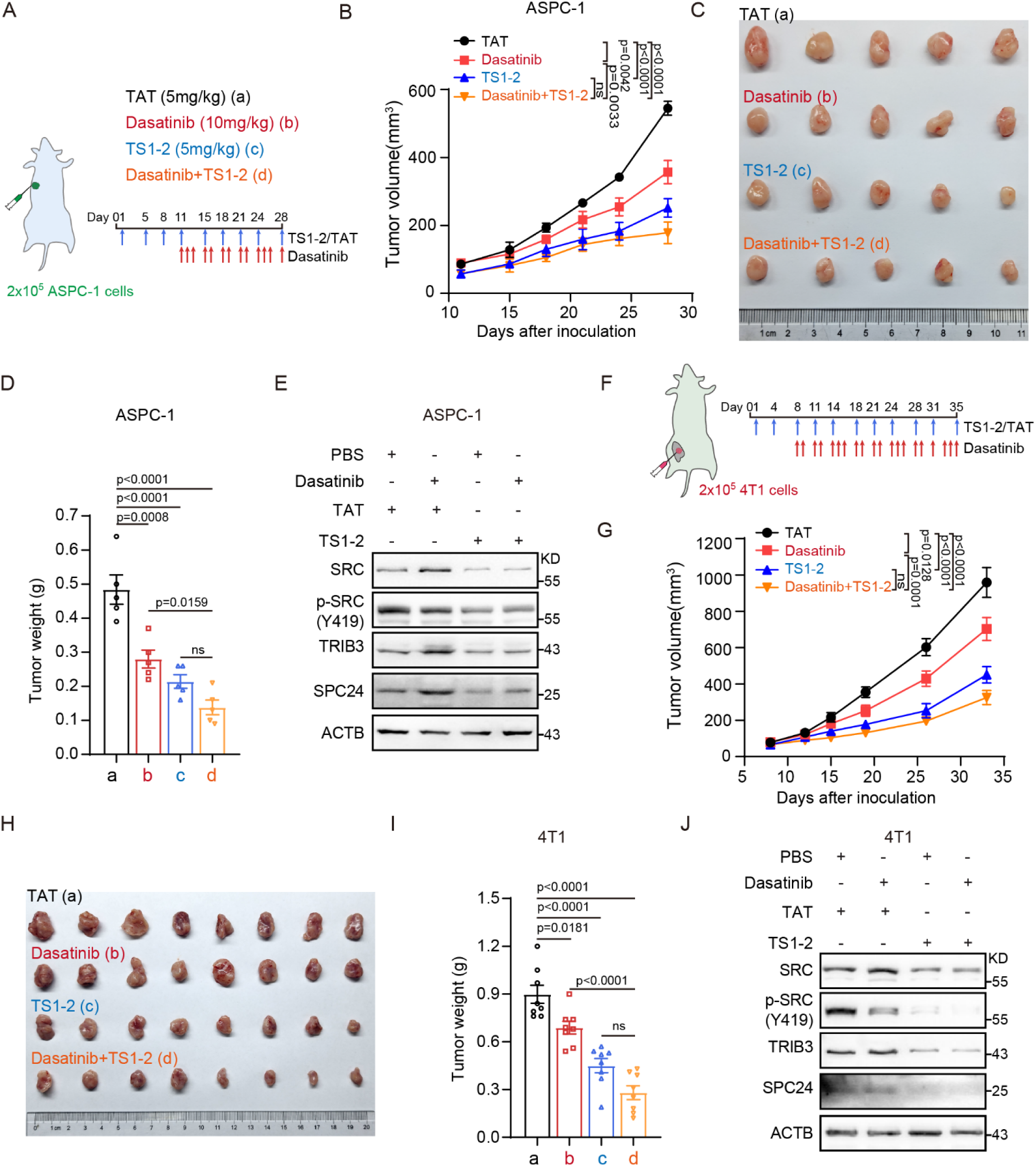
TS1-2 synergistically reduced ASPC-1 and 4T1 xenografted tumor progression with Dasatinib. **A,** Schematic diagrams of the ASPC-1 xenografted BALB/c nude mice model and the treatment strategies. **B-D,** TS1-2 synergistically reduced the ASPC-1 tumor volume (B-C) and tumor weight (D) with Dasatinib. **E,** The effect of TS1-2 and Dasatinib on the levels of the indicated proteins. Primary ASPC-1 tumor cells were isolated from peptide-treated engrafted mice. Cell extracts were prepared, and the levels of the indicated proteins were detected by immunoblotting. **F,** Schematic diagrams of the 4T1 xenograft BALB/c mice model and the treatment strategies. **G-I,** TS1-2 synergistically reduced the 4T1 tumor volume (G-H) and tumor weight (I) with Dasatinib. **I,** The effect of TS1-2 and Dasatinib on the levels of the indicated proteins. Primary 4T1 tumor cells were isolated from peptide-treated engrafted mice. Cell extracts were prepared, and the levels of the indicated proteins were detected by immunoblotting. n=5 in A-D, n=8 in F-I, n ≥ 3 independent experiments from different animals in E and J. Data are shown as mean ± SEM; *P*>0.05 is considered not significant (N.S.), \**P*<0.05, \*\**P*<0.01, \*\*\**P*<0.001.

## DISCUSSION

The non-catalytic function of kinases govern diverse cellular processes, contribute to kinase inhibitor resistance, and present new opportunities for development kinase-targeted therapeutics (1,16). Nonetheless, only a limited number of small-molecule or peptide modulators targeting non-catalytic functions have been characterized to date. As a non-receptor tyrosine kinase acting as a central signaling hub in multiple oncogenic pathways, SRC plays a pivotal role in the development of human hematologic malignancies as well as a broad spectrum of solid tumors. Currently, clinically approved SRC kinase inhibitors are limited to the treatment of hematologic malignancies, whereas their clinical trials in solid tumors have failed due to ineffectiveness or toxic side effects. Given the complexity of the SRC signaling pathways, resistance to SRC-targeted kinase inhibitors is mainly achieved through the compensatory reactivation of downstream signaling cascades. However, the precise non-catalytic function of SRC and their contributions to drug resistance remains unclear.

In the present study, we discovered that continuous treatment with SRC kinase inhibitors led to prominent nuclear accumulation of SRC, which we hypothesized to be closely associated with its non-catalytic activities. By screening for SRC-interacting proteins enriched in the nucleus following SRC kinase inhibitor treatment, we revealed that the oncoprotein TRIB3 binds SRC and promotes the nuclear accumulation by blocking CHIP-mediated ubiquitin-proteasomal degradation of SRC. Pseudokinase Tribbles 3 (TRIB3, also known as NIPK, or SIKP3) has emerged as a key stress sensor responsive to diverse tumor microenvironmental signals, and frequently acts as a scaffolding partner to induce or sustain chronic inflammation, promote tumorigenesis by impairing homeostatic autophagic flux for oncogenic proteins turnover, and enhance tumor-initiating capacity (51–53). Notably, nuclear-localized, catalytically inactive SRC increased the transcription of both *TRIB3* and *SPC24* by direct promoter binding, ultimately establishing a positive-feedback loop. The abrogation of phosphorylation by SRC kinase inhibitors triggers two parallel oncogenic events. On one hand, inactivation of p-SRC relieves p-STAT1-mediated transcriptional repression of *TRIB3*, and the resulting upregulated TRIB3 interacts with the Nc-SRC to inhibit CHIP-mediated ubiquitination-proteasomal degradation of both TRIB3 and SRC. On the other hand, inhibition of SRC phosphorylation signaling impairs its myristylation and promotes nuclear enrichment of Nc-SRC, which in turn promotes the transcription of oncogenic genes including *TRIB3* and *SPC24* by direct promoter occupancy. These observations indicate that Nc-SRC acts as a transcriptional activator or a co-activator, and thereby potentiates the transcription of target genes. Interestingly, the fused polypeptide TS1-2, designed based on the SRC domain responsible for TRIB3 interaction, blocks the SRC/TRIB3 complex and accelerates the simultaneously proteasomal degradation of both TRIB3 and SRC. This dual degradation consequently suppresses the proliferation, migration, stemness, and xenograft tumor growth of multiple solid tumor cells. Collectively, this study elucidates the mechanism by which the non-catalytic functions of SRC drive resistance to SRC kinase inhibitors, and provides the fusion polypeptide TS1-2 as a promising candidate that impedes solid tumor progression through the coordinated degradation of TRIB3 and SRC.

Acquired resistance to SRC kinase inhibitors that develops during clinical treatment is mainly achieved driven through the compensatory reactivation of downstream signaling pathways cascades. Sustained incubation with Dasatinib results in the restoration of STAT3 activity, which is dependent on the interaction between non-phosphorylated JAK and STAT3 in HNSCC cells (49) and non-small cell lung cancer (NSCLC) cells (50). This finding indicates that non-phosphorylated JAK can still mediate the activation of p-STAT3 through its scaffolding functions. Notably, in contrast to Dasatinib, which inhibits p-STAT3 activity, silencing SRC paradoxically activates p-STAT3 in HNSCC cells (49). Collectively, these results suggest that SRC exerts the multifaceted functions, which are not limited to phosphorylation-mediated activation of downstream cascades. In the current study, we found that Nc-SRC was enriched in the nuclei of surviving cells after following Dasatinib treatment, an effect attributed to the inhibition of SRC myristoylation. Previous studies have reported that non-myristoylated SRC exhibited reduced kinase activity but had enhanced stability compared to myristoylated SRC (54,55). Here, we found that inhibition of the kinase activity of SRC suppressed its myristoylation. These results indicate a potential positive feedback loop linking SRC myristoylation to its kinase activity, consistent with the increased stability of Nc-SRC upon Dasatinib exposure. Currently, the nuclear functions of SRC are mainly exerted through its interaction with and phosphorylation of transcriptional activators or repressors (56,57). Interestingly, our results revealed that Nc-SRC acts as a transcriptional activator by directly binding to the promoters of *SPC24* and *TRIB3* and upregulating their transcription. Silencing Nc-SRC reduced the cell proliferation, migration and sphere formation in Dasatinib-resistant cells, while p-SRC was significantly suppressed in ASPC-1/BMS-R and HUH7/BMS-R cells. Collectively, although the mechanism underlying the inhibition of SRC myristoylation subsequent to the suppression of SRC phosphorylation by SRC kinase inhibitors, as well as the critical DNA binding domain of SRC that mediates its binding to the *TRIB3* and *SPC24* promoters are still need further investigation. These results indicated that the non-catalytic functions of SRC constitute a key factor contributing to resistance to SRC kinase inhibitors.

To further dissect the mechanism underlying the nuclear enrichment of Nc-SRC, we performed SRC Co-IP coupled with mass spectrometry in Dasatinib-treated surviving cells and identified a specific interaction between SRC and TRIB3. Interestingly, the E3 ligase CHIP targets both proteins for ubiquitin-proteasomal degradation, and the SRC/TRIB3 interaction mutually blocks their binding to CHIP, thereby reciprocally stabilizing SRC and TRIB3. The E3 ligase FBXL7 mediates the proteasomal degradation of active SRC through Ser104 phosphorylation and subsequent ubiquitination (58). Therefore, inhibition of SRC phosphorylation by the SRC kinase inhibitors stabilizes Nc-SRC, and the robust nuclear accumulation of Nc-SRC consequently promotes the transcription of oncogenic targets. In our previous studies, TRIB3 was found to inhibit the degradation of MYC via UBE3B in lymphoma and EGFR via WWP1 in NSCLC through direct protein-protein interaction, without altering its own expression (59,60). In the present study, the SRC/TRIB3 interaction inhibits CHIP-mediated ubiquitination and proteasomal degradation of both partners in solid tumor cells, by blocking the critical CHIP-binding interfaces on TRIB3 and SRC. CHIP serves as the primary E3 ligase mediating TRIB3 ubiquitination-proteasomal degradation, and Nc-SRC further protects TRIB3 from degradation. Encouragingly, the fused polypeptide TS1-2 designed based the TRIB3-interacting domain of SRC blocked the TRIB3/SRC interaction and accelerated the concomitant degradation of both TRIB3 and SRC, ultimately inhibiting solid tumors progression. Small-molecule-induced targeted protein degradation has emerged as a transformative therapeutic strategy. Proteolysis-targeting chimeras (PROTACs) have been extensively developed to degrade numerous clinically relevant targets, including kinases (61), nuclear receptors (62) and epigenetic enzymes (63). For instance, Dr. Lee’s group constructed an identified a dual IGF-1R/SRC PROTAC by conjugating IGF-1R and SRC inhibitor warheads to the CRBN E3 ligase ligand via diverse linkers. Dr. Soellner’s team reported a potent and selective dual CSK/SRC PROTAC degrader using Dasatinib linked to E3 ligase ligands (64,65). However, the clinical translation of these PROTACs is hindered by suboptimal pharmacokinetic profiles, particularly poor aqueous solubility and limited cellular permeability (66). Notably, the fused polypeptide TS1-2 acts as a PROTAC-like dual TRIB3/SRC degrader for TRIB3 and SRC, with favorable water solubility and cell-penetrating properties. To date, no selective TRIB3 degrader have been documented. Our study therefore provides a promising TRIB3-targeting degrader with potent efficacy for the treatment of solid tumors. With further advanced chemical modifications aimed at prolonging its half-life, this inhibitory polypeptide is expected to exhibit substantially enhanced therapeutic potential. In this study, we uncovered the multifaceted regulatory roles of SRC for cancer development. Phosphorylated SRC (p-SRC) inhibits *TRIB3* transcription through phosphorylated STAT1 (p-STAT1), whereas Nc-SRC acts as a transcriptional activator by directly binding to the *TRIB3* promoter. TRIB3 has been extensively documented to repress gene transcription via interactions with various transcription factors, including ATF4 (67), TCF4 and β-catenin (68), DDIT3 (69), C/EBPβ (70) and PPARγ (71). TRIB3 represses *STAT1* transcription by activating EGFR-STAT3 signaling, which is achieved through stabilizing EGFR via direct protein-protein interaction (60,72). In addition, SRC has been reported to mediate STAT1 activation in PDGF-stimulated NIH-3T3 cells (73). The current study further demonstrated that STAT1 directly binds to the *TRIB3* promoter and represses its transcription, while p-SRC promotes the phosphorylation and activation of STAT1, thereby resulting p-SRC-mediated inhibition of *TRIB3* transcription. Together, these interactions form a TRIB3-STAT1 negative feedback loop. These results elucidate the molecular mechanism underlying the upregulation of TRIB3 following p-SRC inhibition by SRC kinase inhibitors, and identify this regulatory axis as a crucial contributor to the development of resistance to SRC kinase-targeted therapy.

In summary, our study indicates that pseudokinase partner TRIB3 interacts with SRC in contributes to the pathogenesis of multiple solid tumors via its non-catalytic functions, which is coordinated with elevated transcriptional of the oncogenes *TRIB3* and *SPC24*, triggered by activation of the TRIB3-Nc-SRC axis in SRC kinase inhibitor-resistant solid tumors. Our work further elucidates the implications of these findings, and offers potential therapeutic strategies via PPI blockers. Thus, this work provides a proof-of-concept for directly targeting the non-catalytic regions of SRC as a potential combination strategy to overcome SRC kinase inhibitor resistance in solid cancer, achieved through inhibition of the TRIB3/SRC interaction.

## Acknowledgments

This work was supported by grants from the National Natural Science Foundation of China (82073892, 82173853, 82373914, 82574462 and 82525067), the CAMS Innovation Fund for Medical Sciences (CIFMS, 2024-I2M-TS-014, 2021-I2M-1-026), and Fundamental and Interdisciplinary Disciplines Breakthrough Plan of the Ministry of Education of China (JYB2025XDXM609).

## Author contribution

Bing Cui conceptualized and participated in the study’s overall design, supervision, and coordination. Jinmei Yu, Xusheng Wang and Pengyue Liu designed and performed most of the experiments. Zhenhe Wang, Jianli Liu, Xinchao Ban, Qianwen Zhang and Zewen Zhang participated in molecular and cellular biological experiments. Tian Gan, Ning Xu, Jichao Zhou, Jiaojiao Yu and Hongyu Yuan participated in the animal studies. Pingping Li and Xiaowei Zhang provided various advice on experimental design and manuscript writing. Bing Cui, Jinmei Yu, and Xusheng Wang wrote the manuscript. All authors read and approved the manuscript.

## Additional information

During manuscript preparation, the authors utilized the AI tool Doubao for linguistic refinement, including polishing writing fluency and conducting grammar and spelling proofreading to eliminate textual errors.

