## Supplementary methods, Figure and Table for "Targeting Non-Catalytic Sites of SRC Sensitizes the Efficacy of SRC Kinase Inhibitors in Solid Tumors"

by Jin-mei Yu et al

### SUPPLEMENTARY MATERIALS

**Supplementary Methods**

**Supplementary Figure 1**

Non-catalytic SRC (Nc-SRC) is involved in tumor proliferation, migration, stemness and nuclear enrichment. Related to Figure 1.

**Supplementary Figure 2**

SPC24 mediates the functions of Nc-SRC. Related to Figure 1.

**Supplementary Figure 3**

TRIB3/SRC interaction is enriched in nucleus after Dasatinib treatment. Related to Figure 2.

**Supplementary Figure 4**

SRC and TRIB3 synergistically promote the cell proliferation, migration and stemness. Related to Figure 3.

**Supplementary Figure 5**

CHIP mediates SRC ubiquitination and degradation. Related to Figure 4.

**Supplementary Figure 6**

Schematic diagrams of screening key transcription factors mediating SRC-mediated repression of *TRIB3* transcription. Related to Figure 5.

**Supplementary Figure 7**

TS1-2 inhibits proliferation and stemness of MCF7 breast cancer cells. Related to Figure 7.

**Supplementary Table 1**

SRC protein, mRNA expression and OS/RFS/DFS Prognostic Risk Analysis, Related to Supplementary Figure 1.

**Supplementary Table 2**

TRIB3 OS/DFS Prognostic Risk Analysis, Related to Supplementary Figure 3

**Supplementary Table 3**

Primer Sequences for RT-qPCR. Related to Figure 1, 2, 4 and 5.

**Supplementary Table 4**

Primer Sequences for ChIP-RT-qPCR. Related to Figure 1 and 5.

**Supplementary Table 5**

Primer Sequences for siRNA. Related to Figure 3, 4, 5, 7 and Supplementary Figure 2, 4, 5, 6.

**Supplementary Table 6**

Primer Sequences for plasmid construction. Related to Figure 1, 5, 6, 7 and Supplementary Figure 1, 2, 5.

### Supplementary Methods

#### Cell culture

Human embryonic kidney cell line HEK293T, human liver hepatocellular carcinoma (LIHC) cell line HCCLM3, MHCC97H and HUH7, human kidney renal clear cell carcinoma (KIRC) cell line 769-P and 786-O, human breast cancer (BRCA) cell lines MDA-MB-231, MCF7 and murine breast cancer cell line 4T1 were obtained from National Infrastructure of Cell Line Resource, Peking Union Medical College (Beijing, China) and authenticated via short tandem repeats (STR) profiling. Human pancreatic ductal adenocarcinoma (PDAC) cell lines ASPC-1 and PANC-1 were acquired from American Type Culture Collection (ATCC, Manassas, VA, USA） and validated with the same STR for authentication. All cell lines were cultured under standard laboratory conditions in accordance with the instructions provided by ATCC. mycoplasma contamination was excluded in all cell lines using the MycoAlertTM Mycoplasma Detection Kit (Lonza, Basel, Switzerland; Cat No. LT07-418). None of the cell lines utilized in this study are listed in the database of curated by the International Cell Line Authentication Committee (ICLAC).

#### Plasmid construction

Various SRC truncation mutants were generated and constructed into the pCDH−Hyg-myc expression vector via the BamHI and XbaI restriction sites. The constructed SRC variants included: full-length (FL) SRC (amino acids [aa] 1–536), truncation mutants M1 (aa 260–536), M2 (aa 143–260), M3 (aa 1–143), M4 (aa 306–536), M5 (aa 346-536), M6 (aa 384–536), M7 (aa 260–502), M8 (aa 260-469), M9 (aa 260-439), as well as point mutants $SRC^{Y419F}$, $SRC^{Y419D}$. Additionally, the promoter fragments of *SPC24* and *BIRC5* were amplified and constructed into the pGL3-basic vector via KpnI and XhoI restriction enzyme digestion and ligation.

#### Immunoblot Analysis and Protein Stability Assay.

To assess the stability of the existing proteins, cycloheximide (CHX; MedChemExpress, HY-12320) was employed as a specific protein synthesis inhibitor. For protein stability  evaluation, cells were treated with

20 μM CHX for the specified time periods. Subsequently, cell lysates were prepared in RIPA lysis buffer supplemented with protease inhibitors (Lablead, C0101) on ice, and protein concentrations were quantified using BCA Protein Assay Kits (TIANGEN, China, PA115) according to the manufacturer's instructions. Immunoblot analysis were performed to detect protein expression levels. Proteins (20–60 μg per lane) were separated by sodium dodecyl sulfate-polyacrylamide gel electrophoresis (SDS-PAGE) and subsequently transferred to polyvinylidene difluoride membrane (PVDF) (Bio-rad, USA,1620177). The membranes were incubated overnight at 4℃ with the following primary antibodies: anti-SRC (CST, 2109), anti-p-SRC (Y419) (CST, 6943, The rabbit monoclonal antibody against Phospho-Src Family (Tyr416) was generated with a synthetic phosphopeptide surrounding Tyr419 of human SRC), anti-p-SRC (Y530) (CST, 2105), anti-CHIP (CST, 2080), anti-Myc tag (CST, 2276), anti-Stat1(CST, 9172), anti-p-Stat1(CST, 9167), anti-TRIB3 antibody (Abcam, ab75846), anti-SPC24 (Proteintech, 26268), anti-Flag tag (Proteintech, 20543), anti-GFP tag (Proteintech, 50430), anti-His-tag (Proteintech, 66005), and anti-GAPDH (Proteintech, 60004). All primary antibodies were used at 1:1000 dilution, except for anti-GAPDH antibodies, which was diluted at 1:2000.

**Cytoplasmic and Nuclear Fractionation**

Cytoplasmic and nuclear fractions were isolated using a Cytoplasmic/Nuclear Separation Kit (Solarbio Life Sciences, EX2650) following the manufacturer's instructions. Briefly, HUH7 cells were harvested, lysed in Extraction Buffer A on ice for 30 min, and centrifuged to obtain the cytoplasmic supernatant. The resulting nuclear pellets were further lysed in Preservation Buffer B to extract nuclear proteins. The levels of the indicated proteins in both cytoplasm and nuclear were analyzed by immunoblotting.

**Co-Immunoprecipitation (IP)**

Cell pellets were lysed in Co-IP cell lysis buffer (Beyotime, China, P0013) supplemented with protease inhibitors (Selleckchem, B14001). For immunoprecipitation, lysates were incubated overnight at 4 °C with either anti-Myc (Selleckchem, USA, B26302) or anti-HA (Selleckchem, USA, B26202), or with anti-TRIB3 antibodies (Abcam, 75846). Lysates incubated with primary antibodies were further incubated with Protein A/G Plus-Agarose (Santa Cruz Biotechnology, sc-2003) for 2 h at 4°C with rotation. Protein interaction complexes were eluted by boiling the beads for 10 min, then subjected to SDS-PAGE and analyzed with immunoblotting.

**Mass Spectrometry (MS) analysis**

ASPC-1 cells were treated with dimethyl sulfoxide (DMSO) or Dasatinib for 24 h. Whole-cell lysates were harvested and incubated overnight at 4°C with gentle rotation for immunoprecipitation using anti-SRC antibody (CST, 2109) or anti-IgG isotype control antibody (CST, 2729). The antibody–lysate complexes were then incubated with Protein A/G Plus-Agarose (Santa Cruz Biotechnology, sc-2003) for an additional 2 h at 4°C. Protein interaction complexes were eluted by heating the beads at 98°C for 10 min. The eluted protein samples were then submitted to Beijing Qinglian Biotech, Co., Ltd. for mass spectrometry data acquisition and analysis. The generated mass spectrometry data have been deposited to iProX (Integrated Proteome Resources, https://www.iprox.cn) with dataset identifier (Project ID: IPX0016935000, ProteomeXchange ID:PXD077860 ). The dataset is currently is private with temporary access (Link: https://www.iprox.cn/page/PSV023.html;?url=1777538723056LDAO; code: sT5l)

**Ubiquitination Assay**

*In vivo* ubiquitination assays were performed as previously described (1). Briefly, HEK293T cells were transfected with plasmids encoding 6×His-ubiquitin, *SRC-Myc* and the indicated constructs for 24 h. Buffer A (0.1 M Tris-HCl, 0.1 M $NaH_2PO_4$, 0.05% Tween-20) was prepared. Cells were pretreated with 20 μM MG132 for 6 h at 37°C before harvesting. One-tenth of the cells were lysed in RIPA buffer supplemented with protease inhibitors (Lablead, C0101) on ice for 30 min. The remaining cells were lysed with urea buffer (Buffer A was supplemented with 6 M urea and 10 mM imidazole, pH 8.0) at room temperature for 1 h. Equal amounts of protein were incubated with Ni-NTA agarose beads (Qiagen) overnight at 4°C with gentle rotation. Beads were washed four times with denaturing wash buffer (Buffer A was supplemented with 6 M urea and 20 mM imidazole, pH 8.0) and once with native wash buffer (Buffer A was supplemented with 20 mM imidazole, pH 8.0). Protein complexes were eluted in elution buffer (500 mM imidazole buffer) for 20 min at room temperature with gentle shaking. Ubiquitinated proteins were separated by SDS-PAGE for immunoblotting analysis.

**Immunofluorescence (IF) Staining**

For cellular immunofluorescence, ASPC-1 or HEK293T cells were cultured on sterile slides in a 24-well plate, then treated with Dasatinib or transfected with indicated constucts for 24 h. Cells were fixed with 4% formaldehyde for 20 min at room temperature, permeabilized with 0.5% Triton X-100 for 15 min, and blocked with 3% BSA for 30 min at room temperature. Cells were then incubated overnight at 4°C with anti-SRC (1:100, CST, 2109) or anti-Myc (1:100, CST, 2276) primary antibody. After three washes with PBS, cells were incubated with Alexa Fluor 488-conjugated goat anti-rabbit secondary antibody (Invitrogen, A11034) for 120 min at room temperature in the dark. Following three washes with PBS, nuclei were counterstained and mounted with ProLong Gold Antifade Mountant with 4,6-diamindino-2-phenylindole (DAPI) (Invitrogen, P36935). Fluorescence images were obtained using an upright microscope (ZEISS or Olympus Microsystems).

For tissue immunofluorescence assays, formalin-fixed, paraffin-embedded human liver (OD-CT-DgLiv04), pancreatic (HPanA060CS02) and breast (OD-CT-RpBre03-004) cancer tissue and non-cancer tissue microarrays were purchased from Shanghai OUTDO Biotech Co., Ltd. (China). The human liver cancer microarray included 41 cases of invasive hepatocellular carcinoma and 23 cases of adjacent normal liver tissue. The human pancreatic microarray included 37 cases of invasive pancreatic adenocarcinoma and 23 cases of adjacent normal pancreatic tissue. The human breast cancer microarray included 30 cases of invasive lobular breast carcinoma and 27 cases of adjacent normal breast tissue. After deparaffinization in xylene and rehydration through a graded ethanol series (100%, 95%, 80%, 70%, 50% of ethanol and water, 3 min each step), antigen-retrieval was performed in citrate buffer (10 mM sodium citrate, pH 6). Slides were permeabilized with 0.5% Triton X-100 for 30 min and blocked with 3% BSA for 30 min at room temperature, then incubated overnight at 4 °C with the anti-TRIB3 (1:200, Abcam, ab137526) and anti-SRC (1:100, ThermoFisher, 60315-1-IG) primary antibody. After three PBS washes, slides were incubated with Alexa Fluor plus 488-conjugated goat anti-mouse (ThermoFisher, A32723) and Alexa Fluor 647-conjugated goat anti-rabbit (ThermoFisher, A21244) secondary antibodies for 120 min at room temperature. Nuclei were counterstained with DAPI after three PBS washes. Fluorescence images were obtained using an upright microscope (Olympus Microsystems).

**Real-Time Quantitative PCR (RT-qPCR) and RNA Interference(RNAi)**

Total RNA was extracted from cultured cells using an RNA extraction kit (ESScience, Shanghai, China, RN001) following the manufacturer's instructions. cDNA synthesis was performed from 0.5~1 μg of total RNA using TransScript II One-Step RT-PCR SuperMix (TransGen Biotech, AQ321-01) at 42 °C for 30 min. RT-qPCR were performed with PerfectStart® Green qPCR SuperMix (TransGen, AQ601-02-V2) on an Applied Biosystems QuantStudio 3 Real-Time PCR System (Thermo Fisher). Gene-specific primers listed in Supplementary Table 3. Relative mRNA expression levels were normalized to GAPDH as an internal control.

Small interfering RNAs (siRNAs) targeting human *TRIB3*, *SRC*, *STUB1* and *SPC24* were synthesized by RiboBio. siRNA transfection was performed using Lipofectamine™ RNAiMAX Transfection Reagent (Thermo Fisher, 13778150) according to the manufacturer’s instructions. Gene silencing efficiency was verified by detecting mRNA levels at 24 h and protein levels at 48 h post-transfection, respectively. The sequences of gene-specific siRNAs are listed in Supplementary Table 5.

**Dual-Luciferase Reporter Assay.**

HEK293T cells were seeded in culture plates 24 h prior to transfection. Cells were co-transfected with pRL

Reporter Vector and either *TRIB3-Luc* (provided by professor Hua), *SPC24-Luc* or *BIRC5-luc* reporter vectors. After 24 h of transfection, cells were harvested and lysed for Photinus pyralis (firefly) and Renilla reniformis (Renilla) luminescent assays by using the Dual-Luciferase® Reporter Assay System according to the manufacturer's instruction (Promega, E1910).

**Chromatin Immunoprecipitation (ChIP)**

ChIP was performed using SimpleChIP® Enzymatic Chromatin IP Kit (Magnetic Beads, CST, 9003) according to the manufacturer's instructions. Briefly, 4 X $10^6$ cells were cross-linked with 1% formaldehyde at room temperature for 10 min, and excess formaldehyde was quenched with glycine as previously described (1). After enzymatic hydrolysis and ultrasonic treatment, the sheared chromatin fragments were immunoprecipitated with anti-IgG, anti-Myc-tag, or anti-Flag-tag antibodies at 4°C. ChIP-enriched DNA was reverse-crosslinked, purified and analyzed by quantitative real-time PCR. Primer sequences specific to the target gene are listed in Supplementary Table 4.

**Tumor cell migration assay**

Tumor cell migration assays were performed using 8.0 μm Transwell (Corning). Transwell membranes were pre-coated with 10 μg/mL fibronectin (MCE, HY-P3160) and air-dried before use. Tumor cells were pretreated under various conditions: transfecttion with indicated siRNAs for 36 h; incubation with 10 μM TAT or TS1-2 for 24 h; or treatment with indicated concentrations of Dasatinib (MCE, BMS-354825) for 24 h. $1 \times 10^5$ ASPC-1 cells, $3 \times 10^4$ 769-P cells or $3 \times 10^4$ HUH7 cells were suspended in 200 μl of serum-reduced medium (0.4% FBS) and seeded into the upper chamber of Transwell. The lower chambers were filled with 750 μl of complete medium. Chambers were incubated for 24 h (ASPC-1), 12 h (769-P), and 24 h (HUH7), respectively. The migrated cells in the lower chamber were stained with crystal violet and quantified as previously described.

**Cell viability assay**

ASPC-1, 769-P or HUH7 cells were seeded at a density of $2 \times 10^3$ cells per well into 96-well microplates and cultured for 20 h. Subsequently, cells were treated with the indicated siRNAs, 10 μM TAT/TS1-2, or indicated concentrations of Dasatinib for 96 or 120 h. Cell viability was evaluated using 3-(4,5-dimethylthiazol-2-yl)-2,5-diphenyl tetrazolium bromide (MTT) solution at 5 mg/ml (KeyGEN BioTECH, KGH3101-1). The optical density (OD) at 570 nm was measured using a microplate reader (Biotek).

**Mammosphere assays.**

ASPC-1, 769-P or HUH7 cells were pretreated with indicated siRNAs for 36 h, 10 μM TAT or TS1-2 for 24 h, or indicated concentrations of Dasatinib for 24 h. For mammosphere formation assays, ASPC-1 cells, 769-P cells or HUH7 cells were seeded at a density of 500 cells per well into 96-well ultralow-attachment plates (Corning, 3474) and cultured in StemXVivo Serum-Free Tumorsphere Media (R&D Systems, CCM012). Mammosphere images were acquired using a phase-contrast microscope (ZEISS Microsystems). After 5-7 days of culture, mamospheres (defined at 50μm diameter for 769-P and HUH7 cells and 30 μm for ASPC-1 cells) were manually counted to assess the self-renewal capacity of tumor cells.

**Animal Studies**

BALB/c-nude mice were purchased from GemPharmatech Co., Ltd (Jiangsu, China) and housed in maximum barrier facilities with individually ventilated cages, sterilized food, and water. BALB/c mice were obtained from Beijing HFK Bioscience Co., Ltd. (Beijing, China) and maintained in the animal facility of the Institute of Materia Medica under specific-pathogen free (SPF) conditions. All animal experiments were approved by the Animal Experimentation Ethics Committee of the Chinese Academy of Medical Sciences, and all procedures were conducted following the guidelines of the Institutional Animal Care and Use Committees (IACUC) of the Chinese Academy of Medical Sciences. All animal procedures were complied with the ARRIVE guidelines(2).

For xenograft implantation, tumor cells were suspended in PBS with 50% Matrigel (354230, Corning) on ice before xenograft. For the PDAC xenograft mouse model, $2x10^5$ ASPC-1 cells were subcutaneously injected into the right flank of 8-week-old male BALB/c-nude mice. For the 4T1 xenograft mouse model, $2x10^5$ 4T1 cells were orthotopically implanted into the 4th mammary fat pad of 8-week-old female BALB/c mice. One day after xenograft, mice were intraperitoneally treated with 5 mg/kg TAT or TS1-2 twice weekly for 4 consecutive weeks. Tumorigenesis was analyzed at 4 or 5 weeks after cell transplantation. Once tumor volumes reached approximately 100 $mm^3$, mice were orally treated with 10 mg/kg Dasatinib on a 5-day-on and 2-day-off schedule for 4 weeks. Tumor volumes were measured twice a week. The two perpendicular diameters of the tumors were measured, and tumor volume was calculated using the formula $V = 1/2 \times \text{length (mm)} \times \text{width (mm)}^2$. The final peptide treatment was administered 1 h before primary tumor dissection. Proteins lysates extracted from primary tumor tissues were subjected to Western blotting with the indicated antibodies for protein expression analysis.

**Statistical analysis**

All data are shown as the mean ± standard error of the mean (SEM), and all the presented data are representative of at least three independent replicates. Two-tailed student's t-test was used for comparisons between two groups. For multiple group comparison, one-way ANOVA analysis of variance followed by Duncan's post hoc test, unless otherwise indicated. All statistical analyses were using GraphPad Prism 9 or SPSS 24.0 (SPSS, Inc.). $P<0.05$ was considered statistically significant.

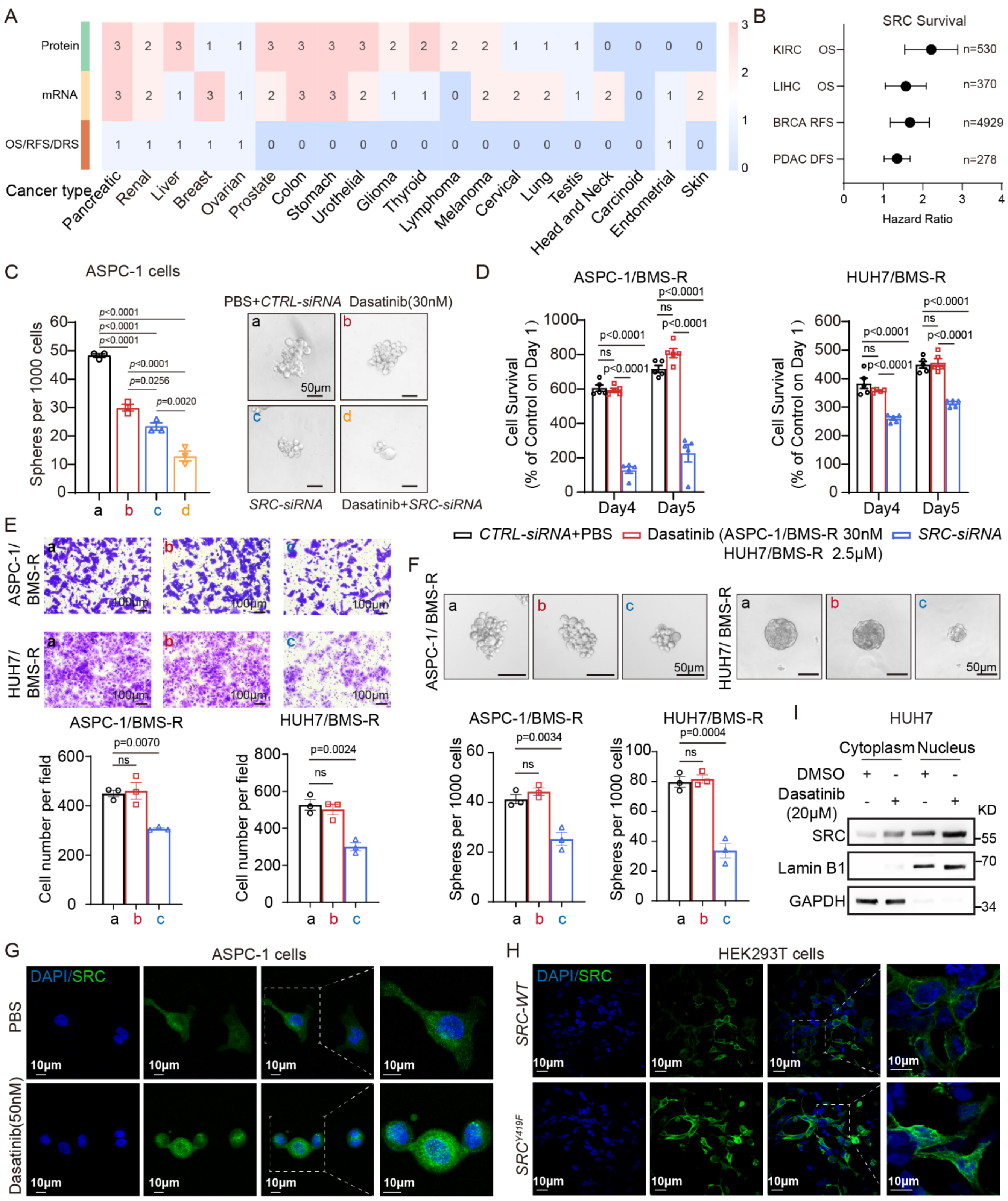


**Supplementary Figure 1.**

**Non-catalytic SRC (Nc-SRC) is involved in tumor proliferation, migration, stemness and nuclear enrichment.** Related to Figure 1. **A**, Heatmap illustrating SRC protein, mRNA expression and OS/RFS/DFS Prognostic Risk Analysis in different cancer types. Protein and mRNA expression were retrieved from HPA database, Kaplan-Meier survival analysis was performed to clarify the prognostic values of SRC across diverse cancers, all data were scale-ranked (detail in supplementary table 1). **B**, Kaplan-Meier survival analysis was performed to clarify the prognostic values of SRC across KIRC, LIHC, PDAC, and BRCA with overall SRC expression values above 4. Overall survival (OS), disease-free survival (DFS), and recurrence-free survival (RFS). **C,** Tumor sphere formation analysis in ASPC-1 cells transfected with *CTRL-siRNA*, *SRC-siRNA* and treated with or without 30nM Dasatinib*.* **D-F,** Tumor proliferation (D), migration (E) and sphere formation (F) analysis in ASPC-1/BMS-R and HUH7/BMS-R cells transfected with *CTRL-siRNA*, *SRC-siRNA* or treated with Dasatinib*.* **G-H,** Immunofluorescence indicates SRC (green) expression in ASPC-1 cells treated with or without Dasatinib (G), or in HEK293T cells transfected with an $SRC^{WT}$-expressing vector or $SRC^{Y419F}$-expressing vector (H). **I,** Western blotting results indicate SRC expression in the cytoplasm and nucleus of HUH7 cells treated with or without Dasatinib treatment. Data represent at least three separate experiments and are shown as the mean ± SEM., $P>0.05$ is considered not significant (N.S.), *$P<0.05$, **$P<0.01$, ***$P<0.001$.

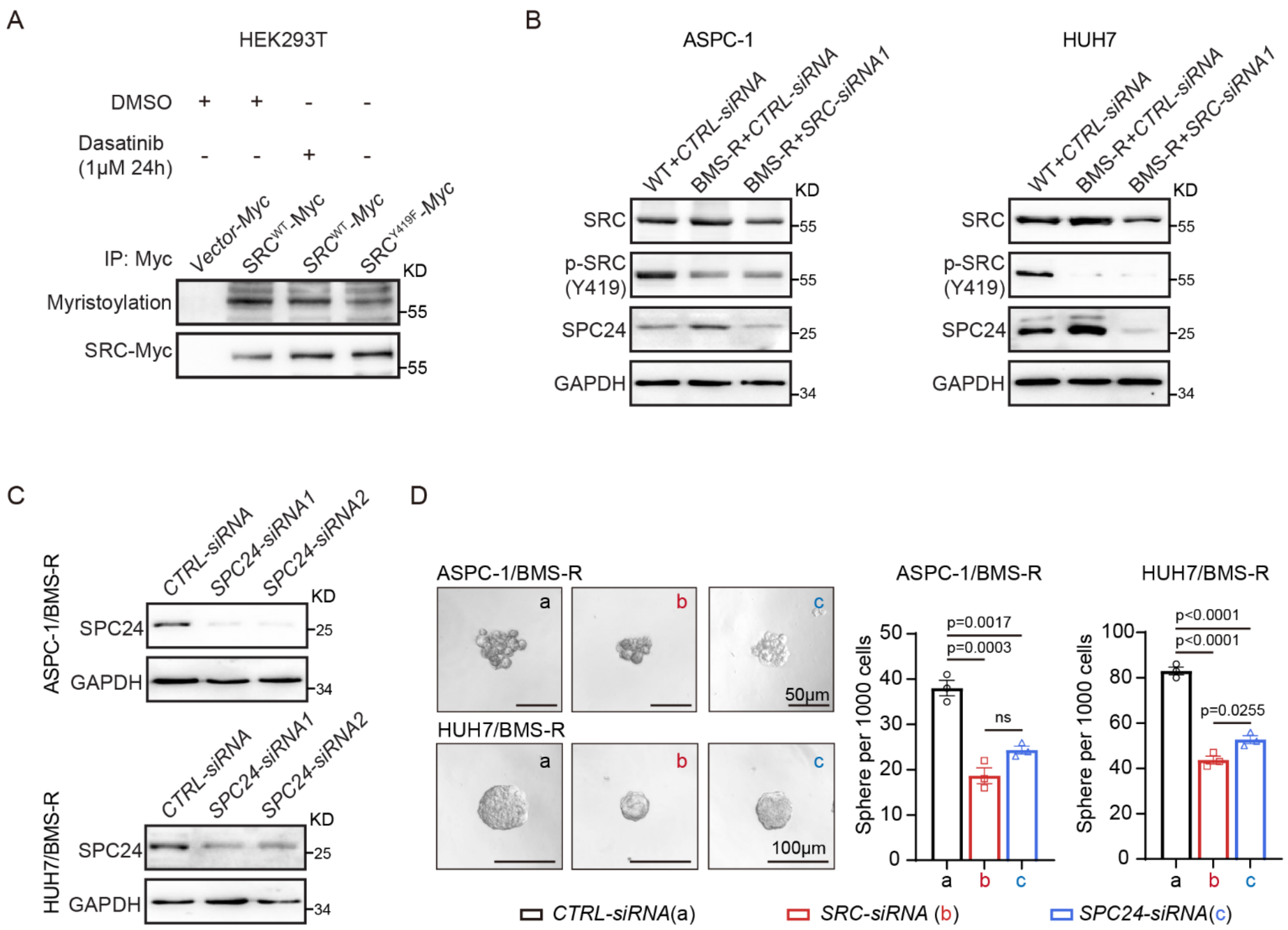


**Supplementary Figure 2.**

**SPC24 mediates the functions of Nc-SRC**. Related to Figure 1. **A,** Kinase-dead *SRC*$^{Y419F}$ decreased myristoylation of SRC. HEK293T cells were transfected with indicated plasmid for 24 h, the cell extracts were IP with anti-SRC-Myc antibody and blotted with anti-SRC-Myc or anti-myristoylation antibody. **B,** Representative Immunoblots probed for SRC, p-SRC (Y419), SPC24 or GAPDH of lysates prepared from ASPC-1, ASPC-1/BMS-R, HUH7 or HUH7/BMS-R cells transfected with *CTRL-siRNA*, or *SRC-siRNA1*. **C,** The representative immunoblots of protein lysates of SPC24 or GAPDH, as indicated on the left, in ASPC-1/BMS-R, HUH7/BMS-R cells transfected with *CTRL-siRNA*, *SPC24-siRNA1* or *SPC24-siRNA2*. **D,** Tumor sphere formation analysis in ASPC-1/BMS-R, and HUH7/BMS-R cells transfected with *CTRL-*

*siRNA*, *SRC-siRNA* or *SPC24-siRNA*. Data represent at least three separate experiments and are shown as the mean ± SEM., *P*>0.05 is considered not significant (N.S.), **P*<0.05, ***P*<0.01, ****P*<0.001.

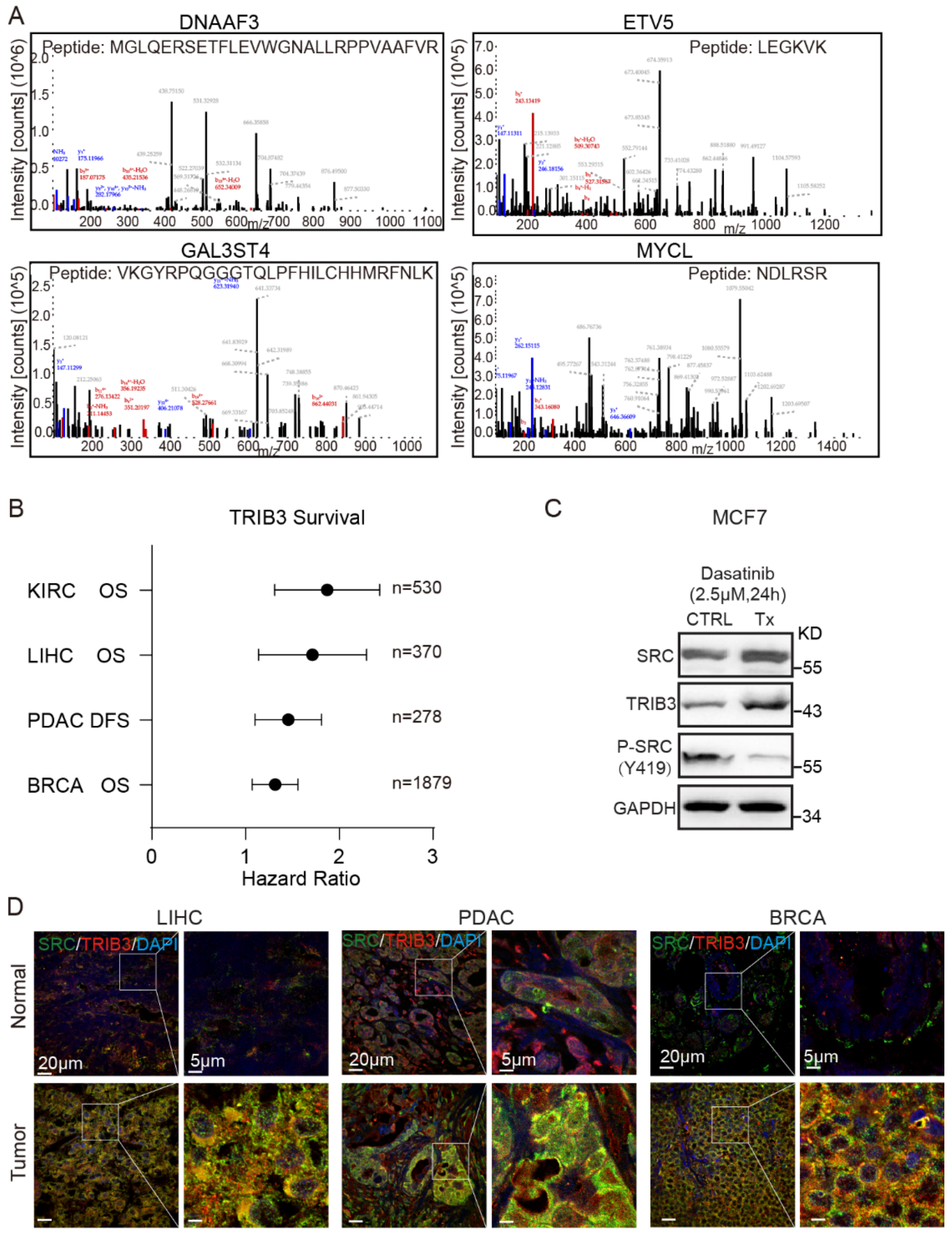


**Supplementary Figure 3.**

**TRIB3/SRC interaction is enriched in nucleus after Dasatinib treatment**. Related to Figure 2. **A,** The Co-IP mass spectrometry data is shown that DNAAF3, ETV5, GAL3ST4 and MYCL were interacted with SRC after Dasatinib treatment for 24 h. **B,** Kaplan-Meier survival analysis was performed to clarify the prognostic values of TRIB3 across KIRC, LIHC, PDAC, and BRCA. Overall survival (OS), disease-free survival (DFS). **C,** Representative Immunoblots probed for SRC, p-SRC (Y419), TRIB3 or GAPDH of lysates prepared from MCF7 cells treated with 2.5 μM Dasatinib for 24 h. **D,** Representative immunofluorescence images for TRIB3 (red) and SRC (green) in human hepatic, pancreatic and breast cancer tissues and adjacent normal tissue. Data represent at least three separate experiments and are shown as the mean ± SEM., *P*>0.05 is considered not significant (N.S.), **P*<0.05, ***P*<0.01, ****P*<0.001.

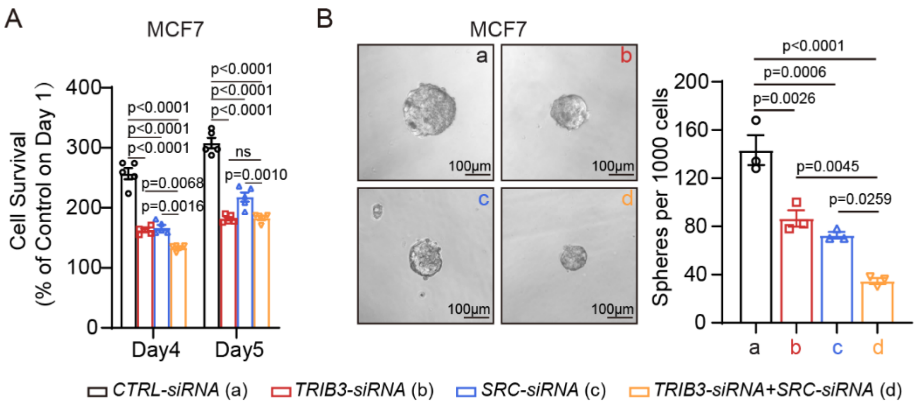


**Supplementary Figure 4.**

**SRC and TRIB3 synergistically promote the cell proliferation, migration and stemness**. Related to Figure 3. Tumor proliferation (**A**) and sphere formation (**B**) analysis in MCF7 cells transfected with *CTRL-siRNA* (a), *TIRB3-siRNA* (b), *SRC-siRNA* (c) or both *TIRB3-siRNA* and *SRC-siRNA* (d). The representative tumor sphere formation photos are presented on the left of each panel (scale bar 100μM). Data represent at least three separate experiments and are shown as the mean ± SEM., P>0.05 is considered not significant (N.S.), *P<0.05, **P<0.01, ***P<0.001.

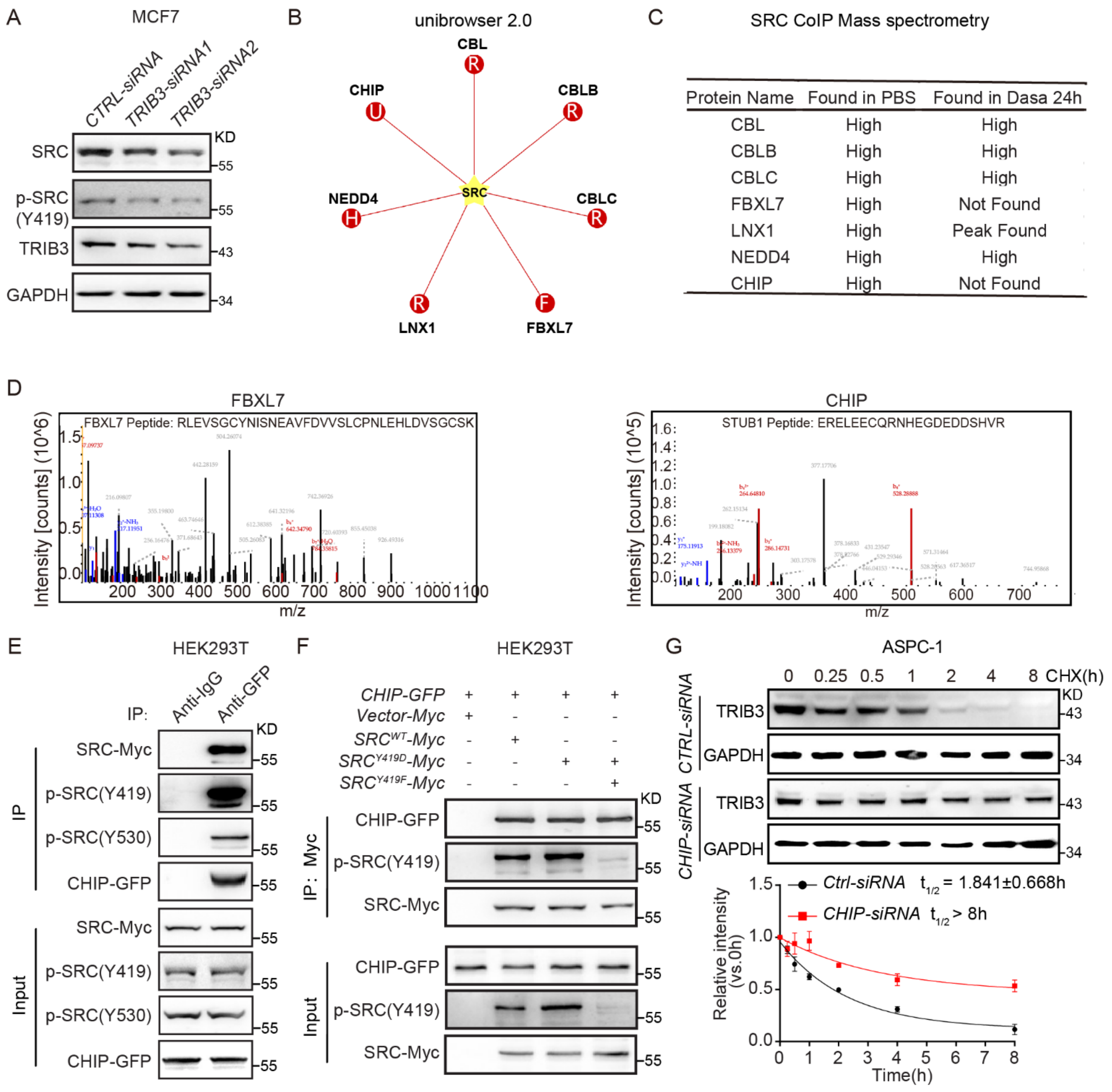


**Supplementary Figure 5.**

**CHIP mediates SRC ubiquitination and degradation.** Related to Figure 4. **A,** Immunoblots of protein lysates, as indicated on the top, transfected with *CTRL-siRNA*, *TRIB3-siRNA1* or *TRIB3-siRNA2* in MCF7 cells. **B,** Predicted ubiquitin ligase (E3)-SRC interactions in eukaryotic species by Unibrowser 2.0 database. **C,** SRC interacted with E3 ligases identified by co-immunoprecipitation (Co-IP) with mass spectrometry in pre-treated or 24 h Dasatinib-treated ASPC-1 cells. **D,** SRC/FBXL7 and SRC/CHIP interactions were only identified by Co-IP with mass spectrometry in pre-treated but not 24 h Dasatinib-treated ASPC-1 cells. **E-F,** CHIP interacts with SRC, p-SRC (Y419), p-SRC (Y530) (E), kinase active *SRC*$^{Y419D}$, and kinase-dead *SRC*$^{Y419F}$ (F). HEK293T cells were transfected with the indicated plasmid for 24 h. The cell extracts were IP with anti-CHIP-GFP antibody (E) or anti-SRC-Myc antibody, and blotted with indicated proteins on the left. **G,** The effect of CHIP (gene name *STUB1*) depletion on TRIB3 degradation *in vitro*. ASPC-1 cells were transfected with *CTRL-siRNA* or *CHIP-siRNA* for 24 h, then treated with CHX (20 μM) for the indicated time. The degradation curves and half-life are indicated on the right panel. Data represent at least three separate experiments and are shown as the mean ± SEM., *P*>0.05 is considered not significant (N.S.), **P*<0.05, ***P*<0.01, ****P*<0.001.

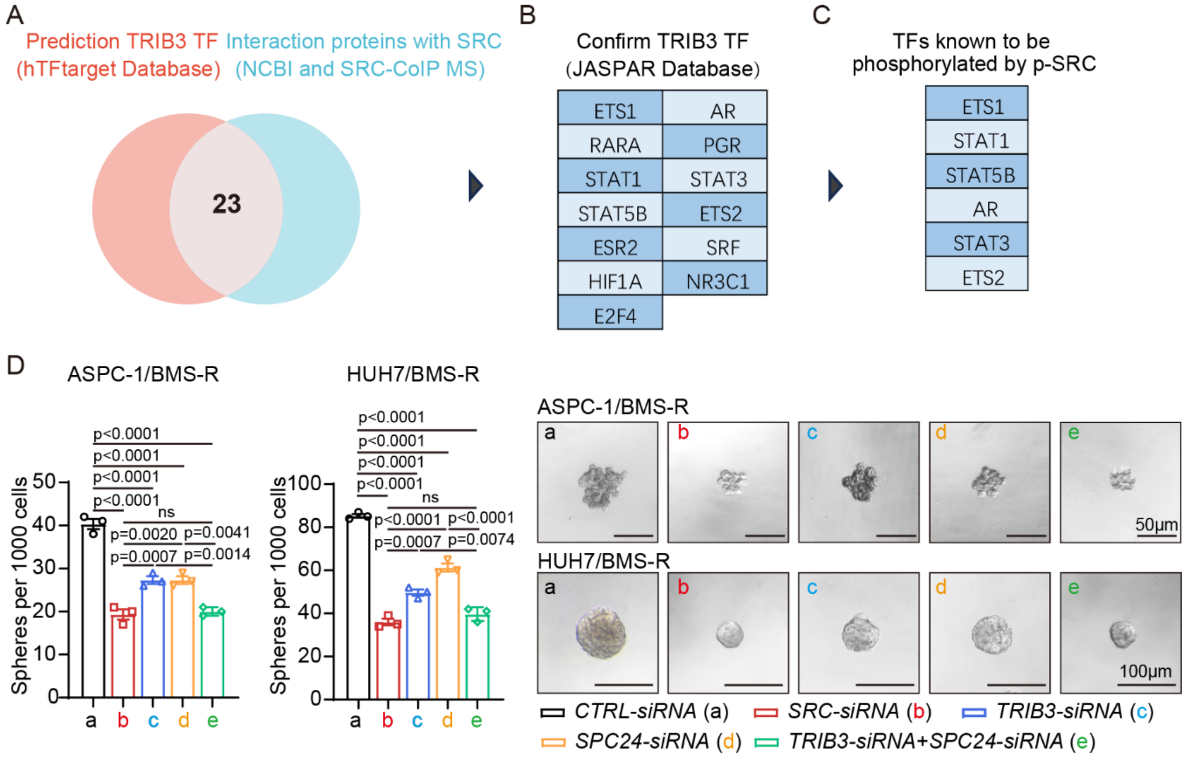


**Supplementary Figure 6.**

**Schematic diagrams of screening key transcription factors mediating SRC-mediated repression of TRIB3 transcription.** Related to Figure 5. **A-C,** Schematic diagrams of screening key transcription factors mediating SRC-TRIB3 transcription loop. First, predicted TRIB3 transcription factors were obtained from hTFtarget database, then 23 common proteins were selected based on the Biomolecular Interaction Network Database (BIND) and SRC Co-IP mass spectrometry data in ASPC-1 cells after 24 h Dasatinib treatment (A). Second, 13 transcription factors were confirmed by JASPAR database (B). Third, 6 transcription factors were selected for their phosphorylation activity by active SRC (C). **D,** Tumor sphere formation analysis in ASPC-1/BMS-R or HUH7/BMS-R cells transfected with *CTRL-siRNA* (a), *SRC-siRNA* (b), *TIRB3-siRNA* (c), *SPC24-siRNA* (d), or both*TIRB3-siRNA* and *SPC24-siRNA* (e). The representative tumor sphere formation photos are presented on the right panel (scale bar 50μm or 100μm). Data represent at least three separate experiments and are shown as the mean ± SEM., P>0.05 is considered not significant (N.S.), *P<0.05, **P<0.01, ***P<0.001.

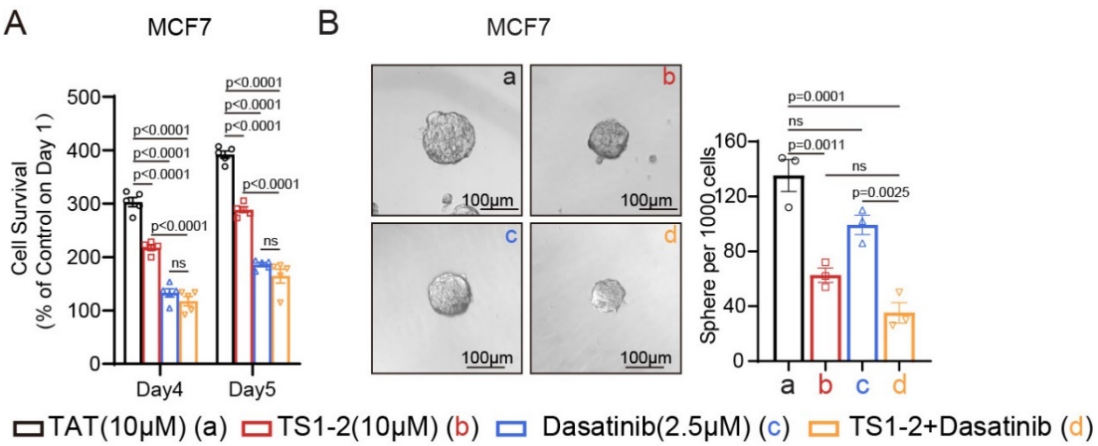


**Supplementary Figure 7.**

**TS1-2 inhibits proliferation and stemness of MCF7 breast cancer cells.** Related to Figure 7. Tumor proliferation (A) and tumor sphere formation (B) analysis in MCF7 cells treated with 10μM TAT (a), 10μM TS1-2 (b), 2.5μM Dasatinib (c) or both TS1-2 and Dasatinib (d). The representative tumor sphere formation photos are presented on the left of each panel (scale bar 100μM). Data represent at least three separate experiments and are shown as the mean ± SEM., *P*>0.05 is considered not significant (N.S.), **P*<0.05, ***P*<0.01, ****P*<0.001

**Supplementary Table 1 SRC protein, mRNA expression and OS/RFS/DFS Prognostic Risk Analysis, Related to Supplementary Figure 1.**

| Cancer type | Protein expression level and score | | RNA expression level and score | | Overall Expression score | OS | | RFS/DFS* | | Risk Score |
|---|---|---|---|---|---|---|---|---|---|---|
| | | | | | | HR | Logrank P | HR | Logrank P | |
| Prostate | ●●● | 3 | ●● | 2 | 5 | 1.13 | 0.8 | 1.28 | 0.24 | 0 |
| Colon | ●●● | 3 | ●●● | 3 | 6 | 1 | 0.97 | 1.09 | 0.48 | 0 |
| Stomach | ●●● | 3 | ●●● | 3 | 6 | 0.79 | 0.15 | 1.46 | 0.26 | 0 |
| Urothelial | ●●● | 3 | ●● | 2 | 5 | 0.62 | 0.0013 | 0.73 | 0.38 | 0 |
| Liver | ●●● | 3 | ● | 1 | 4 | 1.48 | 0.026 | 1.49 | 0.017 | 1 |
| Pancreatic | ●●● | 3 | ●●● | 3 | 6 | 1.33 | 0.18 | 1.31* | 0.0361* | 1 |
| Glioma | ●● | 2 | ● | 1 | 3 | ND | ND | ND | ND | ND |
| Thyroid | ●● | 2 | ● | 1 | 3 | 0.53 | 0.21 | 1.9 | 0.11 | 0 |
| Renal | ●● | 2 | ●● | 2 | 4 | 2.11 | <0.001 | 1.15 | 0.79 | 1 |
| Lymphoma | ●● | 2 | ND | ND | ND | ND | ND | ND | ND | ND |
| Melanoma | ●● | 2 | ●● | 2 | 4 | ND | ND | ND | ND | ND |
| Cervical | ● | 1 | ●● | 2 | 3 | 0.67 | 0.099 | 1.01 | 0.99 | 0 |
| Ovarian | ● | 1 | ● | 1 | 2 | 1.1 | 0.47 | 1.28[#] | 0.011 | 1 |
| Breast | ● | 1 | ●●● | 3 | 4 | 1.28 | 0.13 | 1.29[&] | <0.001 | 1 |
| Lung | ● | 1 | ●● | 2 | 3 | 1.03 | 0.86 | 1.2 | 0.4 | 0 |
| Testis | ● | 1 | ● | 1 | 2 | ND | ND | ND | ND | ND |
| Head and Neck | ND | ND | ●● | 2 | ND | 1.11 | 0.44 | 0.58 | 0.16 | 0 |

| | | | | | | | | | | |
|---|---|---|---|---|---|---|---|---|---|---|
| Carcinoid | ND | ND | ND | ND | ND | ND | ND | ND | ND | 0 |
| Endometrial | ND | ND | ● | 1 | ND | 1.56 | 0.037 | 1.13 | 0.64 | 1 |
| Skin | ND | ND | ●● | 2 | ND | ND | ND | ND | ND | ND |

**Notes：** SRC protein and mRNA expression across 21 solid tumors was analyzed using Human Protein Atlas (HPA) database. TCGA datasets were further interrogated by Kaplan-Meier Plotter to assess the prognostic value of SRC expression. Based on HPA database data, SRC protein expression was classified by the percentage of SRC-positive patients across various cancers: 3 (50%-100%), 2 (20%-50%), 1 (0.08%-20%), ND= not detected (0%); SRC mRNA expression was classified by the median of pTPMs across various cancers:3 (50%–100%), 2 (20%–50%), 1 (1%–20%), and ND (not detected) for 0%.

Kaplan-Meier survival analysis was performed to clarify the prognostic values of SRC across diverse cancers with overall SRC expression values above 4. Patients were categorized into high and low SRC expression subgroups using the median cut-off values, with overall survival (OS), recurrence-free survival (RFS) and disease-free survival (DFS*) selected as survival evaluation indicators. Risk score 1 was given when SRC showed risk predictive potential in any single survival endpoint including OS, RFS and DFS*.

[#]Ovarian (SRC probe:1558211_s_at), [&]Breast (SRC probe:213324_at).

**Supplementary Table 2 TRIB3 OS/DFS Prognostic Risk Analysis, Related to Supplementary Figure 3**

| Cancer type | OS | | | DFS | | |
|---|---|---|---|---|---|---|
| | HR | Logrank P | Probe | HR | Logrank P | Probe |
| Liver | 1.61 | 0.0066 | | 1.2 | 0.27 | |
| Pancreatic | 0.94 | 0.78 | | 1.41 | 0.00707 | |
| Breast | 1.29 | 0.0078 | 218145_at | 1.239 | 0.35 | |
| Renal | 1.78 KIRC | 0.00021 | | 0.93 | 0. 9 | |

**Notes：** Survival analysis was performed using the Kaplan-Meier Plotter (KM plot) online database to explore the association between TRIB3 expression and patient survival across various cancer types. Patients were split by the median value of TRIB3 expression into high and low expression groups. The survival endpoints analyzed included overall survival (OS), recurrence-free survival (RFS), and disease-free survival (DFS).

**Supplementary Table 3 Primer Sequences for RT-qPCR. Related to Figure 1, 2, 4 and 5.**

| Human gene name | Primer sequences | |
|---|---|---|
| | Forward | Reverse |
| *TRIB3* | GCTTTGTCTTCGCTGACCGTGA | CTGAGTATCTCAGGTCCCACGT |
| *SRC* | ACATCCCCAGCAACTACGTG | TTTCTCGCACGAGGAAGGTC |
| *ZCWPW1* | GATGGCTCAAGAGGCAGAACAG | TGGGCTGTTCAAACCAGAGAGC |
| *UPK2* | CACTGAGTCCAGCAGAGAGATC | ACAGAGAGCAGCACCGTGATGA |
| *TSNAXIP1* | GCCAACTTTGGAGATGTGGTCC | GGATCTCATGCTCCTTGCGAAC |
| *TPT1* | TTGGACTACCGTGAGGATGGTG | CAGAAGCCAGTTATGATGACAGG |
| *SYTL2* | TGTGTTGAGCCTGAGCCATCTC | TGGAAGGCATTTTCCTAGCGGC |
| *SPC24* | GGGATTATGAGTGTGAGCCAGG | ACTCCAGAGGTAGTCGCTGATG |
| *SOCS3* | CATCTCTGTCGGAAGACCGTCA | GCATCGTACTGGTCCAGGAACT |
| *SLC11A1* | CATCCTCACGTTCACCAGCATG | CCACGAAGTAGAGGTTGATGGC |
| *SAPCD2* | TGCTGAAGGAGCAGAACCGACT | TGAAGGTGGAATCCAGAGGTCC |
| *S100A11* | CCAGAAGTATGCTGGAAAGGATG | CATCATGCGGTCAAGGACACCA |
| *RASGEF1C* | CCAAGGGACTTCCAGGAAGAGT | AGCTTCTGGTGCAGAGCCTGTA |
| *RASD1* | CACCGCAAGTTCTACTCCATCC | GGTTGTCCAGACTGAACACCAG |
| *GIT1* | CCGAGAGTTTGCCACCTTGATC | ACGCTGTCGTAGTCGTGTTGGT |
| *GADD45A* | CTGGAGGAAGTGCTCAGCAAAG | AGAGCCACATCTCTGTCGTCGT |
| *EEF2* | CCTCTACCTGAAGCCAATCCAG | CCGTCTTCACCAGGAACTGGTC |
| *CCDC159* | GAAAACGCAGGTGAAATGCCGC | CACTCAGTTCCTTCTGGAGGTC |
| *C1QTNF6* | CTGCCTGAGATCAGACCCTACA | TTGTCACCCTTGCTGCCCTGAG |
| *BNIPL* | CCAAGTTCCACCTCTAAGCTGG | GCCGAAGCAGTGCCAGAAATGC |
| *BIRC5* | GATGACGACCCCATGCAAAG | CGCACTTTCTCCGCAGTTTC |
| *ROHF* | AAGACCTCGCTGCTCATGGTGT | GTGTCGTAGAGGTTCAGGGTCA |
| *CCDC57* | GCTCCTAAACCTGCTGGTGACA | GCTCCAAAACCTCCAGGTGTAC |
| *LRRC27* | CAGGAAAGTACCACTGAATCCGC | TTGCTCACGCATTTGGAGGACG |
| *ACAP1* | CCTGACTCAGAAAGGCGGTTCT | CATCAAGGCGAGCCTGACTGAA |
| *KCNN1* | TGCTGGTCTTCAGCATCTCCTC | CGTAGCCAATGGAGAGGAAGGT |

| | | |
|---|---|---|
| *DNAAF3* | GTCCACTTCCTGCCGCTCAATT | CTCAGGGATGAGAAGATGGACC |
| *ETV5* | GTGTTGTGCCTGAGAGACTGGA | CGACCTGTCCAGGCAATGAAGT |
| *GAL3ST4* | CCTTCCACATCCTCTGTCACCA | TAGTAGGAGAAGGCAGAGCGAG |
| *MYCL* | AAGCGACTCGGGTAAGGACCTC | TTCAGCCAAAGAGAGGCGGAGA |
| *GAPDH* | GTCAAGGCTGAGAACGGGAA | TCGCCCCACTTGATTTTGGA |

**Supplementary Table 4 Primer Sequences for ChIP-RT-qPCR. Related to Figure 1 and 5.**

| Human gene name | Primer sequences | |
|---|---|---|
| | Forward | Reverse |
| *TRIB3-1* | GGTGGAGTGCACTCTTAGG | CCTAAGGTCTGCTGAGCTTCC |
| *TRIB3-2* | TCTGAACCACTTGCCCATAGC | GAGAAGTGGACATGCGGGAAT |
| *TRIB3-3* | CAAGTAGCACCAAGCCCTCC | CGGATGCTCAGTGGAGTGTAA |
| *TRIB3-4* | CTAAGAAAGGTGCTGGACCCAG | GTAGAGACAGGGTTTCTCCATGTTGG |
| *BIRC5-1* | GGGCTGGAGGGCTAATAAGG | GGCATCACATCCACTCACTT |
| *BIRC5-2* | CATTTGTCCTTCATGCCCGTC | GACAGGAGAGCTTTACAGGGT |
| *BIRC5-3* | GCGTTCTTTGAAAGCAGTCGAG | GAGTTGTAGTCCTCCCGCC |
| *BIRC5-4* | GACTTACTGTTGGTGGGACGC | GACTGGGCCACTACCGTGATA |
| *BIRC5-5* | GGAAACAGGCAAAACATAAACAG AAAATCTGG | GTTTCACCATGTTTGCCAGGCTG |
| *SPC24-1* | TGCCATCTTAACATCTTGTTTCTTG | GCTGACTCTAAGGTGCTGGT |
| *SPC24-2* | GTATCCCACGCACACACTGTA | GGTGGGGTTCTAACGGCAAAG |
| *SPC24-3* | TCATCACCGATGACTGACGG | CGCGCAAATTTGACTGTCCG |
| *SPC24-4* | CCAGCTACATGCTAGGATTCTATGTATG | CCACAATCATAGCTCATTGCAACTTCG |
| *SPC24-5* | GGAGGCTGAGGCAGGAGAATG | GTCAGTGCAGACATCAGCCACG |

**Supplementary Table 5 Primer Sequences for siRNA. Related to Figure 3, 4, 5, 7 and Supplementary Figure 2, 4, 5, 6.**

| Human gene name | siRNA sequences | |
|---|---|---|
| | #1 siRNA | #2 siRNA |
| *TRIB3* | CGGTTGGAGTTGGATGACAACTTAG | GATCTCAAGCTGTGTCGCTTT |
| *SRC* | GAGCGGCTCCAGATTGTCAACAACA | CCTACTGCCTCTCAGTGTCTGACTT |
| *SPC24* | CTTTACCACCAAGTTAGTAAA | CTCAACTTTACCACCAAGTTA |
| *STUB1* | GAAGAGGAAGAAGCGAGACAT | GACGCATTCATCTCTGAGAAT |

**Supplementary Table 6 Primer Sequences for plasmid construction. Related to Figure 1, 5, 6, 7 and Supplementary Figure 1, 2, 5.**

| Plasmid name | Primer sequences | |
|---|---|---|
| | Forward | Reverse |
| *SRC$^{WT}$* | CATAGAAGATTCTAGAATGGGTAGCAACAAGAGC | CTTCGCGGCCGCGGATCCAAAATTACAGGTCCTCCTCTGAGATCAGCTTCTGCTCGAGGTTCTCCCCGGGC |
| *SRC$^{Y419D}$* | GAAGACAATGAGTTCACGGCGCGGCAAGG | CGCGCCGTGAACTCATTGTCTTCAATGAG |
| *SRC$^{Y419F}$* | GAAGACAATGAGGACACGGCGCGGCAAGG | CGCGCCGTGTCCTCATTGTCTTCAATGAG |
| *SRC-M1* | CATAGAAGATTCTAGAATGAAGGATGCCTGGGAG | CTTCGCGGCCGCGGATCCAAAATTACAGGTCCTCCTCTGAGATCAGCTTCTGCTCGAGGTTCTCCCCGGGC |
| *SRC-M2* | CATAGAAGATTCTAGAATGGACTCCATCCAGGCTG | CTTCGCGGCCGCGGATCCAAAATTACAGGTCCTCCTCTGAGATCAGCTTCTGCTCGGCCAGGCCCTGAGTC |
| *SRC-M3* | CATAGAAGATTCTAGAATGGGTAGCAACAAGAGC | CTTCGCGGCCGCGGATCCAAAATTACAGGTCCTCCTCTGAGATCAGCTTCTGCTCGGAGGGCGCCACGTAG |
| *SRC-M4* | CATAGAAGATTCTAGAATGTCTCCAGAGGCCTTC | CTTCGCGGCCGCGGATCCAAAATTACAGGTCCTCCTCTGAGATCAGCTTCTGCTCGAGGTTCTCCCCGGGC |
| *SRC-M5* | CATAGAAGATTCTAGAATGAAGGGGAGTTTGCTG | CTTCGCGGCCGCGGATCCAAAATTACAGGTCCTCCTCTGAGATCAGCTTCTGCTCGAGGTTCTCCCCGGGC |
| *SRC-M6* | CATAGAAGATTCTAGAATGAACTACGTCCACCGG | CTTCGCGGCCGCGGATCCAAAATTACAGGTCCTCCTCTGAGATCAGCTTCTGCTCGAGGTTCTCCCCGGGC |
| *SRC-M7* | CATAGAAGATTCTAGAATGAAGGATGCCTGGGAG | CTTCGCGGCCGCGGATCCAAAATTACAGGTCCTCCTCTGAGATCAGCTTCTGCTCCCAGCACTGGCACATG |
| *SRC-M8* | CATAGAAGATTCTAGAATGAAGGATGCCTGGGAG | CTTCGCGGCCGCGGATCCAAAATTACAGGTCCTCCTCTGAGATCAGCTTCTGCTCCATCCCAGGGTAGGGC |
| *SRC-M9* | CATAGAAGATTCTAGAATGAAGGATGCCTGGGAG | CTTCGCGGCCGCGGATCCAAAATTACAGGTCCTCCTCTGAGATCAGCTTCTGCTCATAGAGGGCAGCTTCT |
| *SPC24*-Promoter | CTCTATCGATAGGTACCCAGGCGCGGTGGCTCACG | CTTAGATCGCAGATCTCGAGACTACGCCAGGCTCCAACCCG |
| *BIRC5*-Promoter | CTCTATCGATAGGTACCGGCCGGGCGCAGTGGC | CTTAGATCGCAGATCTCGAGGCCGCCACCTCTGCCAACGG |
